# Bridge recombinase enables versatile rewriting of bacterial genomes

**DOI:** 10.64898/2026.04.29.721476

**Authors:** Jaymin R. Patel, Sophia E. Swartz, Agnès Oromí-Bosch, Liana Yong, Zachary W. LaTurner, Jack E. Demaray, Angela Voelker, Ryuichi Ono, Patrick Vu, Preeti Rao, Hannah Luskin, Pia Andrade, Michael L. Cui, Givi Mchedlishvili, Madeline M. Hayes, Natalie Aluwihare, Carlos E. Iglesias-Aguirre, Emma C. MacKenzie, Cynthia I. Rodriguez, Suzanne Devkota, Spencer Diamond, Brady F. Cress

## Abstract

Bacteria drive crucial processes across ecosystems and impact human health, yet tools to rewrite microbiomes remain limited. Here, we show that bridge recombinase enables versatile and programmable genome editing across the bacterial tree of life. In *Escherichia coli*, we achieved 142 kb insertions at >90% efficiency, megabase-scale inversions (2.3 Mb), and pathway-scale 50 kb excisions. With a single ortholog and bridge RNA (bRNA), we edited bacterial isolates spanning five phyla and diverse members of two human gut communities. We overcame cross-reactivity between co-expressed bRNAs to establish search-and-replace Targetable Recombinase Assisted DNA Exchange (TRADE) editing and demonstrated capture and interphylum transfer of chromosomal pathways, enabling programmable horizontal gene transfer. These advances establish bridge recombinase as a foundation for reprogramming gene flow in complex microbial communities.

## Introduction

Bacteria are among the most abundant and functionally diverse organisms on the planet, driving biogeochemical cycles, bolstering industrial biotechnology, and impacting human health. Most bacteria exist within complex microbial communities whose collective functions emerge from interactions among diverse species, yet our ability to genetically manipulate these organisms, particularly non-model species and intact microbiomes *in situ*, remains limited (*1*–*4*). Advances in genome-scale engineering have demonstrated the power of massive DNA manipulation, including genome minimization (*5*), genetic code reassignment (*6*), and chromosomal restructuring (*7*), but these achievements required iterative construction in a small number of intensively domesticated model organisms. Additionally, structural variation in genomes is increasingly appreciated as a driver of complex microbial ecologies (*8*–*10*), further motivating the development of strategies to precisely interrogate them. The vast majority of prokaryotic diversity, including environmentally and clinically important microbes, remains largely inaccessible (*11*) to the programmable and versatile large-scale genome rearrangements that would enable systematic study and manipulation of these organisms.

Among existing genome engineering tools, DNA recombinases stand out for their ability to mediate diverse, large-scale rearrangements, including insertion, excision, and inversion, in a single step without double-strand breaks or dependence on host repair pathways (*12*–*14*). These properties have made Cre (*15*), Flp, and other recombinases foundational genetic tools and increasingly attractive for genome design and therapeutic applications. However, these classical recombinases are not programmable DNA editors; instead, they recombine around fixed DNA attachment sequences, requiring that their recognition sites be pre-installed at desired genomic loci before rearrangement can occur. This constraint precludes their use at native, unmodified genomic sites. In contrast, the recently characterized bridge recombinases, encoded by IS110-family insertion sequences, combine RNA-guided programmability with the full DNA-editing versatility of recombinases (*16, 17*). In these systems, a single non-coding bridge RNA (bRNA, also referred to as seekRNA) specifies both recombination sites (referred to as donor and target) through two independently programmable loops, each specifying, in the case of *E. coli* variant IS621, a user-defined 14 nucleotide DNA sequence containing a conserved core dinucleotide (CT) (**Fig. 1A**). Reprogramming these bRNA loops enables scarless insertions, inversions, and excisions at user-defined sequences without pre-engineering of the target genome. Bridge recombinases have been shown to mediate programmable DNA recombination in *E. coli* and, through the recent discovery of the ISCro4 ortholog, in human cells (*18, 19*). Several key frontiers remain unexplored. It is unknown how broadly these systems function and can be programmed across bacterial species and in microbiomes. Additionally, the intrinsic biochemical challenges associated with multi-bRNA editing have not been resolved, precluding certain classes of DNA manipulations.

**Figure 1.**
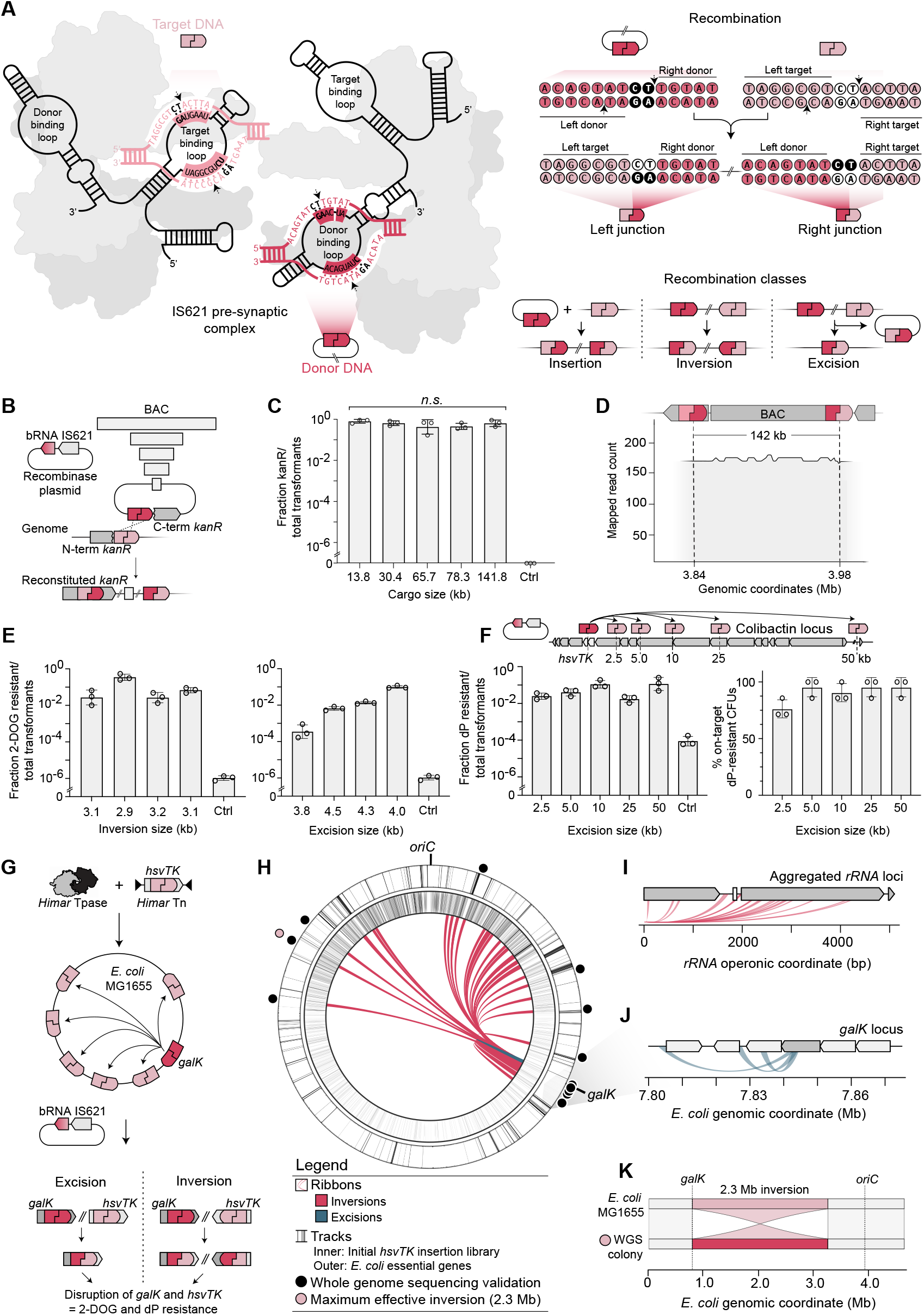
Versatile genome-scale manipulations with bridge recombinase. (**A**) Schematic of IS621-mediated recombination and possible recombination classes with single bRNAs. Nucleotide-resolution of the donor DNA (dDNA) and target DNA (tDNA) substrates during an “insertion”-style recombination. (**B**) Schematic of the split kanamycin assay, wherein a split kanamycin gene is used to assess the recombinase-mediated integration of different-sized plasmids into a programmed locus. (**C**) Fraction of colonies surviving kanamycin selection over total transformants is shown. We observed that insertion efficiency did not statistically vary with insertion size (one-way ANOVA, p-value = 0.35). (**D**) An alignment of whole genome sequencing reads from a representative 141.8 kb recombinant colony is mapped across the *in silico* predicted map. (**E**) bRNAs are programmed to invert or excise the *galK* locus. Following transformation of pEdit, recombinants (fraction resistant to 2-DOG) are quantified. (**F**) bRNAs are designed to excise increasing portions of the colibactin biosynthetic gene cluster. *hsvTK* is inserted into the left flank of the pathway to mediate quantification. Upon transformation of pEdit, excision efficiencies are quantified as fraction resistant to dP. The fraction of post-selection clones with correct excision are quantified via colony PCR. (**G**) To assess genome-wide rearrangements, the counterselectable *hsvTK* gene was randomly inserted across the *E. coli* MG1655 genome (*52*) via *Himar* transposition; bRNAs are programmed to recombine between the native *galK* locus and *hsvTK* insertion library, the outcome being an excision or inversion based on the orientation of *hsvTK* insertion. Upon recombinase induction, recombined cells are selected on 2-DOG-and dP-containing media. (**H**) 67 colonies were randomly selected for recombination mapping via whole genome sequencing. Within the genome plot, inner ribbons represent observed inversions (red) and excisions (blue); the inner ring shows the distribution of initial *hsvTK* insertions; the outer ring highlights essential genes; external spots note rearranged clones validated though whole genome sequencing. (**I**) *rRNA* operons were collapsed into a single contig to enable mapping of inversions at these identical loci. (**J**) Rearrangements proximal to *galK* are enlarged. (**K**) Representative Mauve alignment of a mutant colony encoding a 2.3 Mb inversion (as verified by whole-genome sequencing) against the reference genome *E. coli* MG1655, verifying on-target inversion and the absence of other structural rearrangements. All measurements of fraction resistant over total transformants or resistant CFUs are the mean ± sd of three biological replicates.

Here we show that the IS621 enables programmable genome restructuring across diverse prokaryotes and microbiomes. In *E. coli*, we achieved programmable insertion efficiencies >90% maintained for payloads approaching 142 kb, scarless inversions spanning half the chromosome (2.3 Mb), and a 50 kb excision of the oncogenic colibactin biosynthetic gene cluster (BGC), approaching the technical limits of insertion, inversion, and excision size in this organism. We demonstrate large programmable edits in bacterial isolates spanning five phyla and achieved metagenomic editing of enriched human gut communities without prior strain isolation. Furthermore, we develop a dual-bRNA strategy to enable homologous recombination (HR)-independent search-and-replace editing, a modality we term Targetable Recombinase Assisted DNA Exchange (TRADE). We expand TRADE to catalyze the capture and interphylum transfer of functional chromosomal loci. Together, these results establish bridge recombinase as a generalizable platform technology for large-scale bacterial genome engineering and controlled manipulation of horizontal gene transfer (HGT) in complex microbial communities.

## Results

### Bridge recombinase performs programmable genome-scale manipulations in *E*. *coli*

Prior work has demonstrated bridge recombinase-mediated editing at individual genetic loci in *E. coli (16, 18)*, but whether efficiency is maintained across increasing cargo sizes and genomic rearrangement distances is untested. To determine the size limits of bridge recombinase-mediated recombination in bacterial genomes, we quantified integration of a panel of Bacterial Artificial Chromosomes (BACs) sized at 13.8, 30.4, 65.7, 78.3, and 141.8 kb into the *E. coli* genome (**table 1**) (*20*). Bridge recombinase was expressed *in trans* (on plasmid pEdit) using a dual transcript architecture, which we found most effective (**fig. S1**). A bRNA was programmed to mediate recombination between two halves of a *kanR* gene (**tables 2, 3**). Using this split kanamycin assay to select for on-target recombinants, we observed efficiencies up to 90.9% without significant change in efficiency across increasing cargo size (**Figs. 1B, C**). Whole genome sequencing confirmed full-length, on-target integration of the programmed payloads (**Fig. 1D; fig. S2**).

We then quantified the efficiency of kilobase-scale excisions and inversions using a galactokinase (*galK*) counterselection assay (*21*), in which edits were programmed to disrupt *galK* in the *E. coli* genome and thereby confer resistance to 2-deoxy-galactose (2-DOG). Counterselection for on-target recombinants showed excision efficiencies up to 10.0% and inversion efficiencies up to 36.4% (**Fig. 1E**). To test excision at pathway scale, we targeted the oncogenic (*22*–*24*) and clinically-relevant 54 kb colibactin BGC in *E. coli* Nissle 1917. For quantification, we flanked the gene cluster with the counterselectable herpes simplex virus thymidine kinase (*hsvTK*) marker (*25, 26*), which would be disrupted upon excision and confer resistance to 6-(β-d-2-deoxyribofuranosyl)-3,4-dihydro-8H-pyrimido [4,5-c][1,2] oxazin-7-one (dP). We designed bRNAs to programmably excise 2.5, 5.0, 10, 25, and 50 kb of the pathway. Between 1.82-14.0% of induced pEdit transformants survived counterselection, the proxy for excision efficiency; 71.4-100.0% of these post-counterselection CFUs were bona fide excisions as verified by cPCR (**Fig. 1F**). Efficiencies did not decrease with increasing excision sizes, supporting the scalability of this approach to large genomic deletions.

This size-independent performance led us to hypothesize that bridge recombinase could drive extensive chromosomal rearrangements across the genome. To test this, we performed unbiased genome-wide recombination experiments in which the bridge recombinase was programmed to recombine a fixed endogenous counterselectable gene (*galK*) with a second counterselectable gene (*hsvTK*) that we randomly distributed around the *E. coli* genome using the *Himar* transposase (**Fig. 1G**). Recombinant mutants were picked from dual counterselection plates, analysed by arbitrarily-primed PCR (AP-PCR) (*27*), and whole-genome sequenced. We recovered mutants with 1119 - 4099 bp excisions (**Fig. 1H; figs. S3, S4; table 4**) and a wide size range of chromosomal inversions spanning up to 2.3 Mb, representing the largest effective inversion possible in *E. coli* at approximately half of its genome size (**Figs. 1I-K**). Interestingly, we observed that inversions at ribosomal *rRNA* operons were enriched compared to the distribution of the initial *hsvTK* library (**Fig. 1I; fig. S5**). Taken together, these results demonstrate that bridge recombinase mediates a versatile set of programmable, scarless genome rearrangements in bacteria, ranging from gene and pathway scale to chromosome scale.

### IS621 functions broadly across bacterial phyla

To establish bridge recombinase as a tool for bacterial genome editing beyond *E. coli*, we assessed whether IS621 could function in phylogenetically diverse bacterial isolates. Homologs of IS621 are prevalent across all major bacterial and archaeal phyla (**fig. S6**), suggesting their recombination mechanism may have minimal reliance on host-specific factors, potentially enabling efficient, broad host-range editing. To test this, we selected eleven diverse Gram-negative and Gram-positive species spanning five phyla (Pseudomonadota, Bacteroidota, Bacillota, Deinococcota, and Actinomycetota) of industrial, agricultural, or therapeutic relevance (**Fig. 2B; table 5**). To facilitate bRNA programming across diverse recipients, we identified a 14mer sequence within the *16S* ribosomal RNA gene (*rRNA*) that is universally conserved across bacteria. We constructed a series of pEdit vectors encoding the *16S*-targeting bRNA, the IS621 bridge recombinase, taxa-specific promoters, and antibiotic selection cassettes. These pEdit plasmids were constructed with both replicative backbones for sustained, persistent expression of bridge recombinase, and non-replicative backbones for transient expression (**Fig. 2A**).

**Figure 2.**
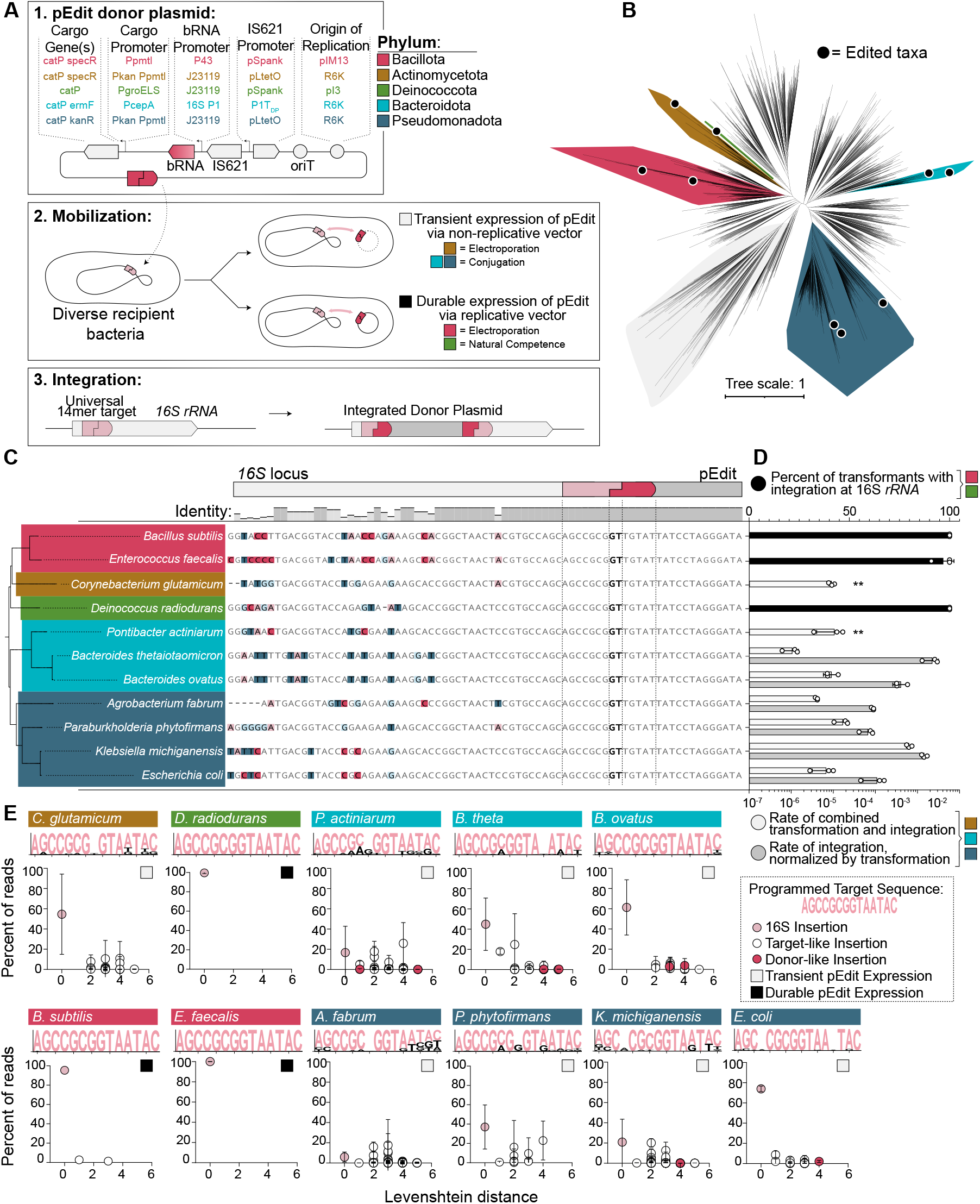
Programmable insertions with bridge recombinase across the bacterial tree of life. Plasmid design and transformation method for phyla-specific pEdit vectors. The bRNA targets a universally-present 14mer in the *16S rRNA* gene. Insertions are quantified via antibiotic selection. (**B**) Unrooted order-level tree derived from GTDB taxa distances (*53*), to illustrate edited strains and phyla. Scale represents relative evolutionary divergence (RED) (**C**) Insertions across diverse species are validated through sequencing of the *16S rRNA* locus. Horizontal phylogenetic distances are derived from GTDB taxa distances (**D**) Top axis: for strains transformed with replicating vectors, the percent of transformant CFUs with PCR-verified integration at 16S, is plotted (black). Bottom axis: for strains transformed with non-replicating vectors, the combined frequency of transient transfer and insertion of the cargo plasmid is quantified (white), as antibiotic-resistant CFUs divided by total recipients. In parallel, a replicative vector is transformed to quantify transformation efficiency, allowing calculation of transformation rate-normalized editing efficiency (black) (mean ± sd of three biological replicates). ** indicates a suitable transformation efficiency control vector was not available for normalization. (**E**) Sequence similarity and off-target profile of bridge recombinase-mediated integration across diverse bacteria. Genomic DNA from transformed populations (> 200 CFUs) was analyzed by AP-PCR to map cargo insertion sites. Insertions were classified as target-like or donor-like and plotted by their Levenshtein distance from the programmed target sequence (x-axis) versus percent of reads (y-axis). Insertions at *16S rRNA* are highlighted in pink; donor-like insertions are shown in red; other target-like insertions are shown as open circles. Sequence logos depict aligned target-like insertion sites for each species, with height weighted by count frequency. Bases matching the programmed target sequence are highlighted in pink.

We observed successful integration of the full 6.4 - 10.3 kb pEdit plasmid at the target site across all 11 targeted species, including *Bacillus subtilis, Enterococcus faecalis, Corynebacterium glutamicum, Deinococcus radiodurans, Pontibacter actiniarum, Bacteroides thetaiotaomicron, Bacteroides ovatus, Agrobacterium fabrum (*previously classified as *Agrobacterium tumefaciens), Paraburkholderia phytofirmans, Klebsiella michiganensis*, and *Escherichia coli* (**Fig. 2C**). When a non-replicative pEdit vector was used, the integration frequency was > 1 x 10^-4^ across tested species, after normalizing for transformation efficiencies using a replicative plasmid control (**Fig. 2D**). In contrast, with replicative pEdit vectors, the fraction of transformant CFUs harboring an integration at *16S* locus approached 100%, as quantified by cPCR (**Fig. 2D; fig. S7**). We mapped integration sites across all edited strains via AP-PCR. Integrations occurred at the *16S rRNA* gene, at off-target sites with sequence similarity to this programmed target, and, rarely, at sequences resembling the donor (**Fig. 2E; figs. S7-9; tables 6, 7**). Comparing these strategies, replicative vectors consistently yielded higher integrant frequencies and substantially lower off-target integration. Together, these results indicate that sustained recombinase expression improves both efficiency and specificity of editing. These findings establish bridge recombinase as a portable, programmable DNA editing tool for diverse bacteria and, given its shared mechanism, suggests that the genome-scale rearrangements demonstrated in *E. coli* are broadly transferable.

### Metagenomic editing with IS621 bridge recombinase

To date, bridge recombinase-mediated genome editing has only been demonstrated in single-species contexts. However, bacteria naturally exist within complex microbial communities that exert collective functions on their environments. Building on our finding that bridge recombinase functions across diverse microbial isolates, we asked whether it could accomplish genome editing across diverse species directly within microbiomes.

We tested editing in two human gut microbial communities derived from an infant stool sample (*28*) and an adult small intestinal mucosal sample. To optimize editing across taxa, we constructed two non-replicative editing vectors targeting Enterobacteriaceae and Bacteroidaceae, respectively encoding the IS621 bridge recombinase, the universal *16S*-targeting bRNA, and cognate antibiotic resistance markers (**Fig. 3A**). To quantify editing within specific taxa and exclude background resistance from the broader community, edited cells were recovered on taxa-selective media with appropriate antibiotics: MacConkey (MC) agar for Enterobacteriaceae and Bacteroides Bile Esculin Gentamicin (BBEG) agar for Bacteroidaceae. Under these conditions, colony formation requires both growth on the selective medium (taxonomic constraint) and integration of the editing plasmid (antibiotic selection). The Enterobacteriaceae-targeting vector was used in both communities, while the Bacteroidaceae-targeting vector was used only in the adult community.

**Figure 3.**
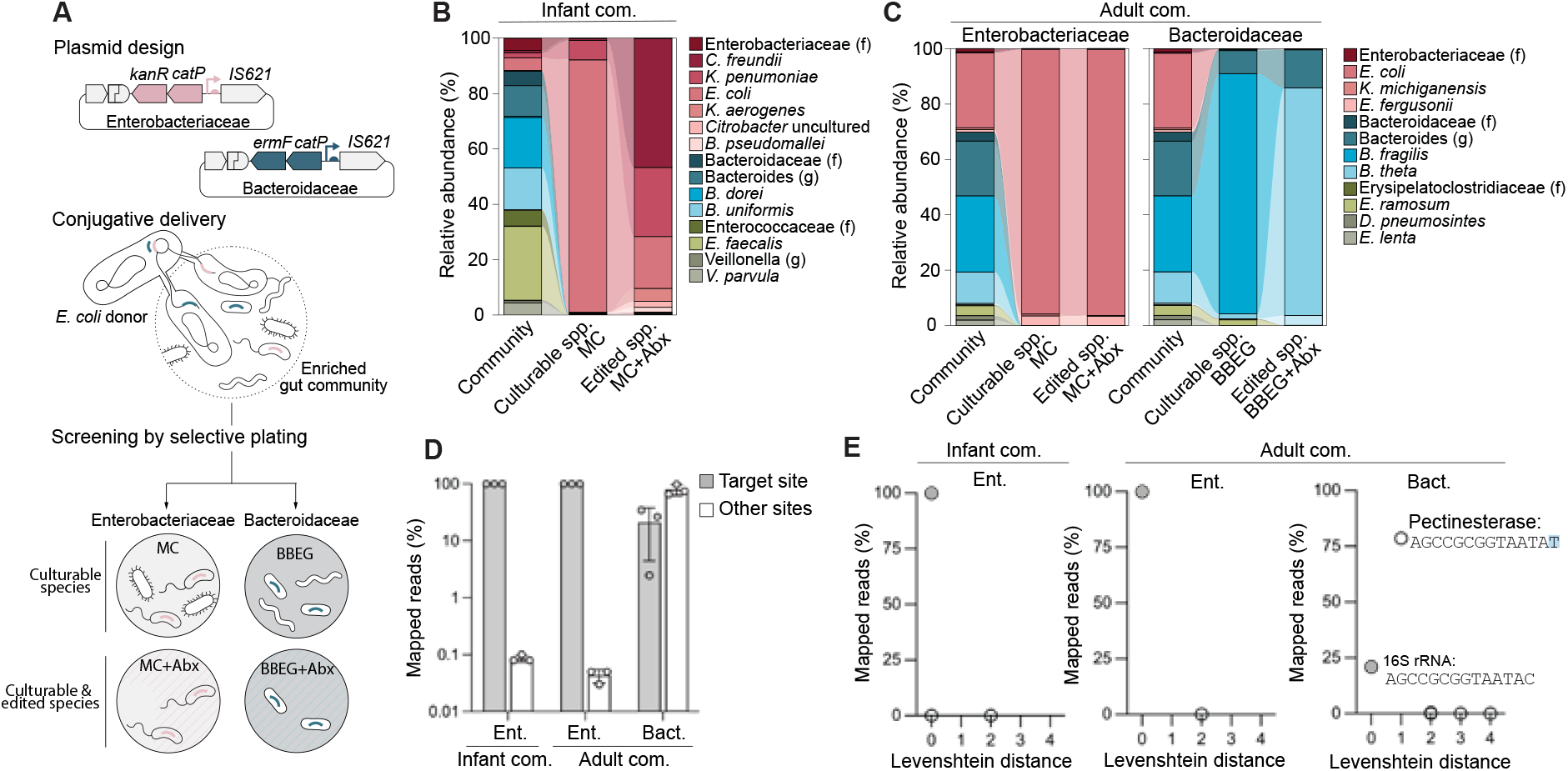
Metagenomic editing with bridge recombinases. (**A**) Schematic of modular bridge recombinase plasmids and workflow for metagenomic editing using bacterial conjugation for editing plasmid delivery. Edited cells are quantified by plating on selective media with and without antibiotics. (**B**) Bacterial composition of infant gut community before conjugation, post-conjugation on selective media, and on selective media with antibiotics (i.e., edited bacteria) as determined over three biological replicates. (**C**) Bacterial composition of adult gut community before conjugation, post-conjugation on selective media, and on selective media with antibiotics (i.e., edited bacteria) as determined over three biological replicates. Left panel on MC media for identification of Enterobacteriaceae and right panel on BBEG for Bacteroidaceae. (**D**) Percentage of IS621 plasmid insertion events at *16S rRNA* locus (on-target) and other loci. (**E**) Levenshtein distances of IS621-mediated target sequences with the fraction of mapped reads. For D-E, mean ± s.d. of three biological replicates are shown. In 3B-C, (f) means family, and (g) genus.

We recovered edited colonies from multiple species in both communities. Editing efficiencies, defined as the fraction of antibiotic-resistant colonies relative to total CFUs on the corresponding taxa-selective medium, reflect the combined efficiency of conjugation and genomic integration across all members of the targeted group. For Enterobacteriaceae, efficiencies were 3.0 × 10^−4^ and 4.7 × 10^−5^ in the infant and adult communities, respectively. For Bacteroidaceae in the adult community, the efficiency was 1.1 × 10^−7^ (**fig. S10**). In the infant community, 16S rRNA sequencing showed that all 11 culturable Enterobacteriaceae species capable of growth on MC were recovered on MC with antibiotics, indicating successful editing across the entire culturable Enterobacteriaceae family (**Fig. 3B**). In the adult community, we recovered edited *E. coli* and *B. thetaiotaomicron* in MC with antibiotics and BBEG and antibiotics, respectively (**Fig. 3C**). Metagenomic insertion mapping via AP-PCR revealed that > 99.9% of Enterobacteriaceae insertions were on-target at the *16S rRNA* locus, whereas *B. thetaiotaomicron* insertions were split between the intended site and a single-mismatch off-target site within a non-essential pectinesterase gene (**Figs. 3D, E**). These findings were corroborated by individual colony screening, which identified 100% on-target insertions in Enterobacteriaceae and 30% on-target insertions in *B. thetaiotaomicron*, in line with observations when editing this species in isolation (**fig. S11**). These results demonstrate that bridge recombinases enable direct, programmable genome editing across multiple species within complex microbial communities without the need for prior isolation.

### Dual bRNA recombination enables search-and-replace TRADE editing

Programmable genome editing tools can mediate insertions and, when combined with site-specific recombinases, even rearrangements, but single-step RNA-guided replacement of any large genomic segment with another has remained challenging. We reasoned that a bridge recombinase guided with two co-expressed bRNAs spanning the boundaries of two distinct genetic loci would enable directed intermolecular DNA exchange, or “search-and-replace editing”, akin to a programmable version of the classic Recombinase Mediated Cassette Exchange (RMCE) (*29*) (**Fig. 4A**). We term this programmable reaction Targetable Recombinase Assisted DNA Exchange (TRADE). To test TRADE, we designed two bRNAs to replace a chromosomal *sfGFP-mRFP* region in *E. coli* with a plasmid-encoded chloramphenicol (*catP)* marker. Double recombination would introduce *catP* into the genome while disrupting both fluorescent reporters (**Fig. 4B**).

**Figure 4.**
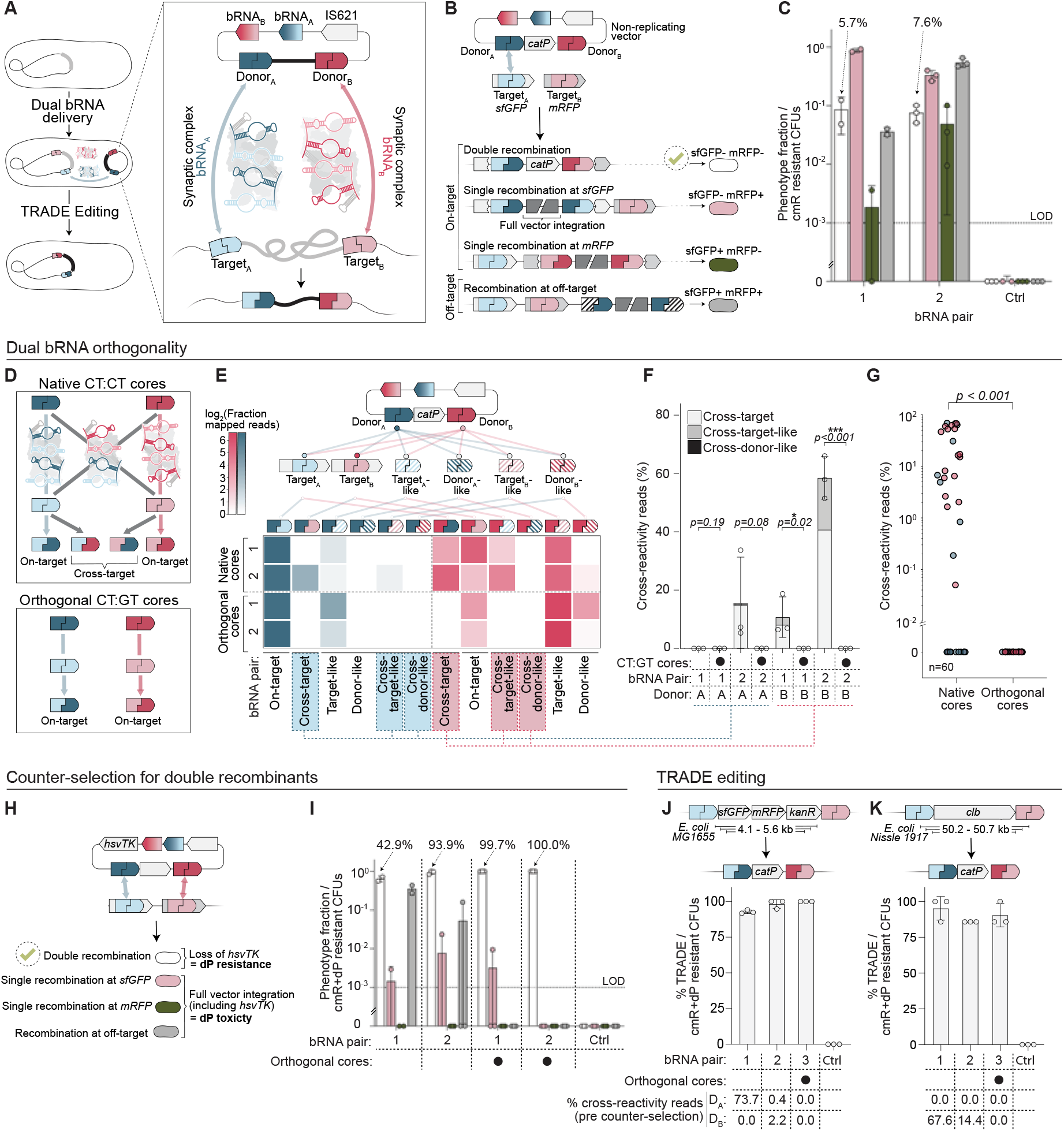
Dual bRNA recombination enables TRADE editing. (**A**) Expression of two bRNAs and formation of two distinct synaptic complexes permit programmable DNA TRADE editing. Dual bRNAs are designed to disrupt *sfGFP* and *mRFP* with a *catP* replacement; incomplete single recombination reactions at *sfGFP, mRFP*, or off-target sites are expected to integrate the full donor plasmid. (**C**) Following recombination, chloramphenicol-selected CFUs are phenotyped by fluorescence to determine recombination type. Double recombinants are the minority product across multiple bRNA pairs. (**D**) Orthogonal dinucleotide core sequences are predicted to suppress cross-reactivity between bRNAs with native CT:CT cores. (**E**) Following transformation and IS621 induction, AP-PCR and deep sequencing are used to map the insertion sites of both donor_A_ and donor_B_. Observed insertion sites are classified by sequence similarity to the true programmed target_A_ and target_B_, or to off-target sites resembling target_A_, target_B_, donor_A_, or donor_B_. Cross-targets are defined as the insertion of donor_A_ into any B-like sequences and vice-versa. The log_2_ fraction of reads from each category is shown as a heat map. (**F**) Cross-target reaction rates are highlighted as a bar graph for each tested bRNA pair (significance via one-tailed t-test). (**G**) Aggregated cross-target reaction rates from 60 independent reactions across 10 bRNA pairs are plotted. Marker colors correspond to donor_A_ and donor_B_ (Welch’s one-tailed t-test, p<0.0001). (**H**) Incomplete single recombination reactions, predicted to integrate the donor plasmid backbone, can be counterselected by placing the conditional toxin *hsvTK* on the backbone. (**I**) Following recombination, dP-selected clones are phenotyped by fluorescence to determine recombination types. (**J-K**) TRADE editing of the *sfGFP-mRFP-kanR* and colibactin BGC using *hsvTK* counterselection and orthogonal cores. The fractions of post-selection CFUs with the correct double recombination are quantified. For all plots, mean ± s.d. of three biological replicates are shown.

Initial experiments using fluorescence phenotyping of chloramphenicol-selected clones revealed that double recombinants were the minority product, comprising 5.7% and 7.6% of recombination products for each bRNA pair (**Fig. 4C; fig. S12**). We identified several failure points limiting double recombination rates. First, we observed that bRNA recombination efficiency varies across bRNAs and target genes, with *mRFP*-targeting bRNAs showing particularly low on-target efficiencies (**figs. S13, S16; table 8**). These results highlight the importance of bRNA design and target selection, which remain poorly understood and difficult to predict.

Second, we identified crosstalk between bRNAs as an additional failure mode. Because each bRNA contains two loops and recombination occurs within a tetramer that has been shown to engage two bRNA molecules (*30*), we wondered if multi-bRNA editing might introduce combinatorial mispairing between non-cognate bRNAs (which we refer to herein as cross-reactivity). If so, this could generate off-target recombination products that scale quadratically with the number of co-expressed bRNAs, in contrast to the linear scaling observed for multiplexed CRISPR systems (note S1). We hypothesized that utilizing orthogonal core dinucleotide sequences (CT:GT) between the two bRNAs could suppress this crosstalk (**Fig. 4D**). Using AP-PCR and deep sequencing to map insertion sites of both donors, and classifying them by sequence similarity to programmed target or donor sites, we found that orthogonal cores eliminated cross-reactivity products (**Figs. 4E-G; fig. S16; table 8**). We observed decreased editing efficiency of CT:GT cores relative to CT:CT cores, consistent with prior work showing lower efficiency recombination at GT relative to CT cores (*16*).

Third, we observed that the majority of recombinants underwent a single recombination event resulting in integration of the full plasmid backbone instead of the intended double recombination event. Incorporating the counter-selectable *hsvTK* marker into the plasmid backbone allowed for negative selection against these single-recombinant intermediates with the nucleoside analog (dP) (**Fig. 4H**). Following dP counter-selection, fluorescence phenotyping confirmed that use of orthogonal cores and counter-selection drove the intended TRADE outcome to nearly 100% (**Fig. 4I**).

Using this optimized strategy, we replaced the ∼4.8 kb *sfGFP-mRFP-kanR* locus in *E. coli* MG1655 and the ∼50.4 kb colibactin biosynthetic gene cluster in *E. coli* Nissle 1917 with *catP*. Across three bRNA pairs for each locus, > 85.7% post-counterselection CFUs exhibited the correct TRADE editing genotype (**Figs. 4J, K; figs. S14, S15**). Here, use of orthogonal cores completely eliminated cross-reactivity products (**fig. S16**). These results establish bridge recombinase-mediated TRADE editing as a powerful method for programmable replacement of pathway-scale genomic segments.

### Dual bRNA recombination enables programmable HGT across phyla

HGT is a major driver of bacterial adaptation and evolution but remains largely unpredictable and difficult to control (*31*). We envisioned that TRADE could be extended to achieve “programmable HGT”: the capture and mobilization of defined chromosomal segments, along with their encoded functions, into diverse hosts (**Figs. 5A, B**).

**Figure 5.**
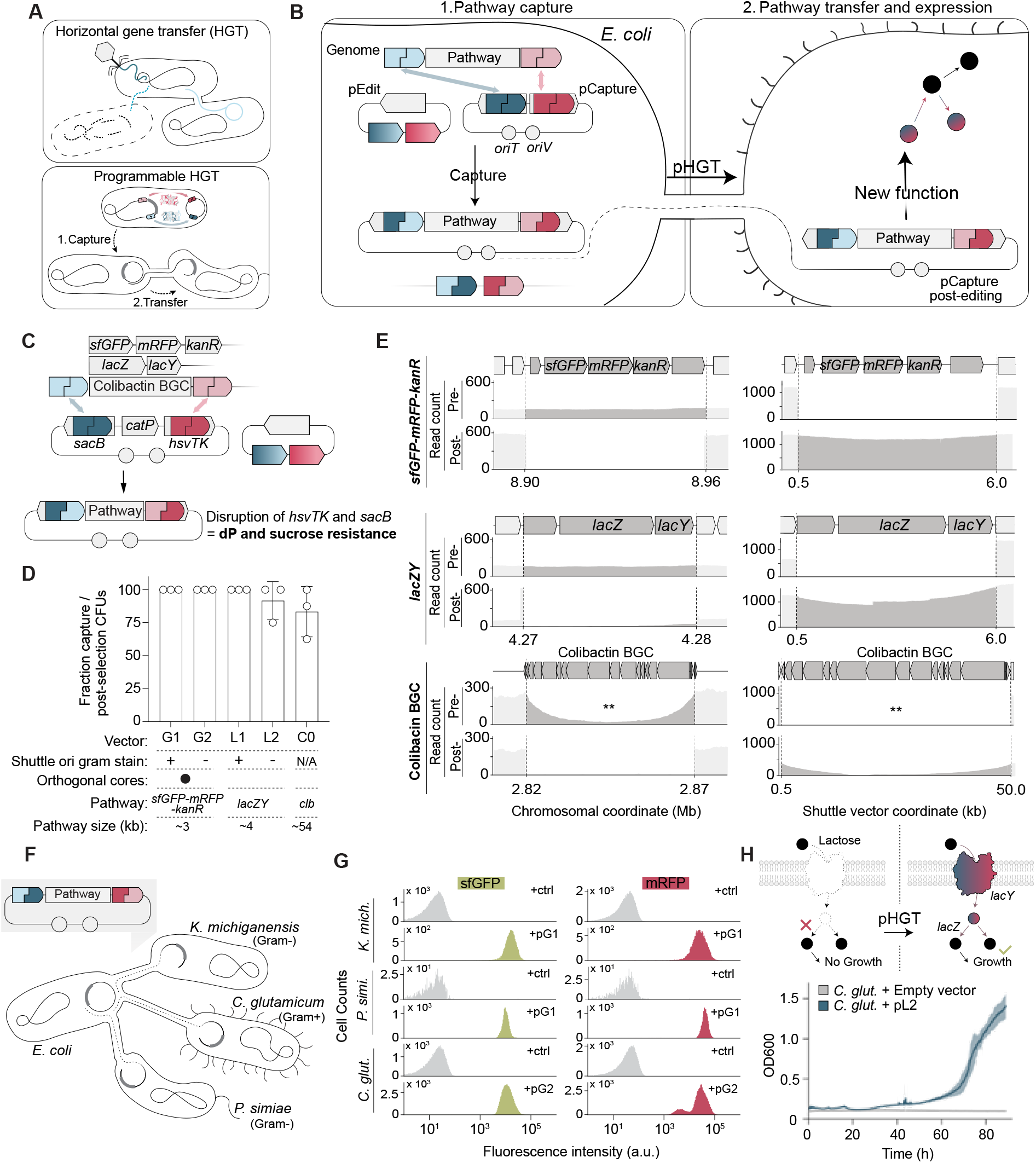
Interphylum programmable HGT with dual bRNAs. (**A**) Using dual bRNAs, we capture defined genomic segments of interest into a shuttle plasmid and transfer them into a species from another phylum. (**B**) Schematic of pathway capture from *E. coli* chromosome into pCapture plasmid, assisted by pEdit plasmid encoding dual bRNAs and IS621 *in trans*. Conjugative transfer of the final pCapture plasmid provides new functionalities to recipient species. (**C**) Three different chromosomal pathways are captured onto vectors encoding counterselectable genes *sacB* and *hsvTK*. Correct pathway capture disrupts both genes and confers dP and sucrose resistance. (**D**) Fraction of *E. coli* CFUs containing the correct pCapture insert after double counter-selection. (**E**) Whole genome sequencing coverage maps confirm the capture of genomic pathways onto pCapture plasmids at the programmed IS621 half sites. **Coverage across the large colibactin pathway drops centrally due to ambiguous read assignment between chromosomally-and plasmid-encoded BGC reference maps. (**F**) Schematic of pathway transfer into *K. michiganensis, P. simiae* and *C. glutamicum* by conjugative delivery. (**G**) Fluorescence intensity of control (+Ctrl, empty vector) and *sfGFP-mRFP-kanR*-expressing strains of *K. michiganensis, P. simiae* and *C. glutamicum* after 36 h of growth in LB media, as measured by flow cytometry using FITC and mCherry filters. (**H**) Growth curves of *C. glutamicum* empty vector control and *lacZY*-expressing strain on RCH2 medium with 60 mM lactose as the sole carbon source confirms activity of the *lacZY* pathway. For all plots, mean ± s.d. of three biological replicates are shown.

To test this, we converted the optimized dual-bRNA editing plasmid into shuttle vectors capable of replicating in phylogenetically distant recipient bacteria and incorporated a dual counter-selection system (**Fig. 5C**). Using these plasmids, we “captured” two chromosomal pathways from *E. coli* with > 83% of correct edits post-selection, including a ∼3 kb fluorescent *sfGFP-mRFP-kanR* and a ∼4 kb lactose-utilizing *lacZY* cassettes. Additionally, we captured a ∼50 kb colibactin BGC, demonstrating the scalability of this technique and its applicability in studying large gene clusters (**Fig. 5D; figs. S17-20**). Whole genome sequencing confirmed complete mobilization of target loci from the chromosome to the plasmid vector (**Fig. 5E**).

Captured pathways were mobilized through conjugation into diverse recipients, including *Klebsiella michiganensis*, an emerging drug-resistant pathogen (*32*), *Pseudomonas simiae*, a plant growth-promoting bacterium (*33*), and the Gram-positive actinobacterium *Corynebacterium glutamicum*, a widely used industrial production organism (*34*) (**Fig. 5F**). Recipient strains stably maintained the transferred plasmids and exhibited the expected phenotypes: *K. michiganensis, P. simiae and C. glutamicum* expressed GFP and RFP fluorescent signal that was readily distinguishible from empty-vector controls (**Fig. 5G; fig. S21**). Programmable HGT of *lacZY* to *C. glutamicum* conferred the ability to grow on lactose as a sole carbon source (**Fig. 5H; fig. S22**), illustrating how programmable HGT can enable access to new nutrients and drive metabolic niche adaptation in microbial ecosystems.

These results demonstrate that bridge recombinases enable HR-independent direct capture and interphylum transfer of functional biosynthetic pathways without prior modification of donor or recipient genomes (*35*), or the requirement for cell-free molecular biology. Together, this work establishes bridge recombinases as a platform for programmable HGT, enabling precise engineering of complex functions across bacterial species.

## Discussion

In this work, we establish bridge recombinase as a single-effector system for programmable insertion, excision, inversion, and replacement of DNA across diverse bacteria and microbial communities. In *E. coli*, bridge recombinase maintained high insertion efficiency for payloads approaching the limits of DNA that can readily be introduced by electroporation, inverted half the chromosome (the largest effective inversion possible for a circular genome), and excised 50 kb of the colibactin biosynthetic pathway, one of the larger contiguous non-essential loci in the *E. coli* genome. Colibactin is a genotoxic metabolite produced by some Enterobacteriaceae and has been implicated in early onset colorectal cancer (*22, 36, 37*), suggesting that programmable excision of its BGC could enable targeted removal of disease-associated small molecules from microbial genomes and communities. Together, these results suggest that bridge recombinase editing scales to the physical and biological limits of the host genome (i.e. transformability constraints for exogenous DNA insertion, distance between essential genes for excisions, and total genome size for inversions), rather than being constrained by the IS621 itself. Beyond *E. coli*, a single bRNA targeting a conserved site in the *16S rRNA* gene directed integration across five phyla, including within enriched human gut bacterial communities, demonstrating that bridge recombinase functions as a broad host-range editor with minimal dependence on host-specific factors. By extending the system to a dual-bRNA modality, we achieved HR-independent single-step search-and-replace TRADE editing, and programmable HGT of chromosomal pathways between phyla; capabilities uniquely simplified by bridge recombinase.

Several important limitations and open questions remain. While bridge recombinase excels at versatile, large-scale edits across diverse genetic backgrounds using a compact two-component system, the specificity of bRNA-directed recombination was variable and difficult to predict from bRNA and genome sequence alone. Off-target insertion rates differed substantially even among closely related species, motivating future systematic investigation of the factors governing bRNA efficiency and specificity. Massive bRNA libraries, combined with machine learning approaches, could reveal how target DNA sequence context, local transcriptional activity, chromosomal topology, and proximity to the origin of replication contribute to recombination outcomes and ultimately enable predictive bRNA design (*38, 39*). For dual-bRNA applications, achieving complete double recombination without counterselection will require a better understanding of how the relative dosage and activity of paired bRNAs govern the balance between single-and double-recombination outcomes (*40, 41*). The observed stalling after the first recombination event may suggest intrinsic autoinhibitory mechanisms, which could mitigate potential toxicity of excessively active IS elements (*42*). Further, eliminating cross-reactivity between co-expressed bRNAs through engineering, directed evolution, or discovery of naturally orthogonal bridge recombinase variants will be equally important. Existing modalities for search-and-replace editing of large DNA segments (*43*) have not been extensively developed for bacteria, perhaps due to the reported challenges of prime editing (*44*) in these microbes (*45*), further motivating the optimization of TRADE editing. Finally, because editing efficiency in non-model organisms is ultimately gated by DNA delivery efficiency, the development of efficient bridge recombinase delivery strategies, including phage-based approaches, will be critical for extending the full capabilities of bridge recombination beyond the species tested here.

It is increasingly clear that microbiome function is determined not only by species composition but also by the metabolic capabilities of community members. These metabolic functions are dynamic and can shift through large-scale genomic changes, including acquisition of biosynthetic gene clusters (*46*), mobilization of genomic islands (*47*), and structural rearrangements (*9*). Together, these processes highlight the need for strategies that enable multimodal editing at comparable genomic scales. Bridge recombinases could enable direct rewriting of genotype and investigation of function across community members *in situ*, including hard-to-cultivate species that grow readily within their native communities (*48*). Future work will test insertions, excisions, inversions, and TRADE editing in diverse bacterial phyla and configure the programmable HGT workflow for one-pot capture, transfer, and integration into the final recipient genome. The ability to capture and transfer defined chromosomal segments between species creates new experimental access to longstanding questions about HGT, including how acquired pathways reshape recipient fitness and how gene flow shapes community function over time (*49, 50*). More broadly, programmable HGT could enable the construction of chimeric microbial genomes engineered within intact communities (*51*), moving beyond observation of natural microbiomes toward their rational redesign. We anticipate that bridge recombinases, together with advances in predictive bRNA design and bacterial delivery systems, will form a foundation for “synthetic microbiomics”, the systematic, programmable orchestration of gene flow within complex microbial ecosystems.

## Acknowledgments

We are grateful to Owen T. Tuck, Siqi Yang, Santiago Lopez, Emily Armbruster, Peter Yoon, and Jérôme Zürcher for critical feedback on this manuscript, to all members of the Cress, Doudna, Rubin, and Diamond labs for helpful discussions, and to Michelle O’Malley, Ben Rubin, Carlotta Ronda, and Yun Song for trainee mentorship. We thank Chloe Summerhill for valuable technical support. We gratefully acknowledge the leadership and support of the Berkeley Initiative for Optimized Microbiome Editing (BIOME), particularly Jennifer Doudna, Jill Banfield, Brad Ringeisen, Audrey Glynn, and Rachel Evans. We thank Phil Fleshner for collecting the healthy adult ileal resection sample used for enrichment of the adult ileal microbiome. We thank Sue Lynch for providing the fecal sample used for enrichment of the infant stool microbiome. We thank Adam Arkin, Adam Deutschbauer, Vivek Mutalik, and Carlotta Ronda for bacterial strains used in the study.

## Author contributions

**Conceptualization: JRP, BFC**

**Experimental Design: JRP, SES, AOB, BFC**

**Experimental Work: JRP, SES, AOB, LY, ZWT, AV, PV, PR, HL, PA, MLC, GM, MMH, NA, CEI, ECM, CIR**

**Computational Analysis: JRP, JED, RO, SD Manuscript Preparation: JRP, SES, AOB, BFC Supervision: SD, SD, BFC**

## Data, code, and materials availability

Sequencing data collected for this study are deposited in NCBI SRA under BioProject accession SUB16150084. Scripts and workflows used for analysis of sequencing runs are available on github under https://github.com/cress-lab/. The insertion_mapper repository contains scripts used to map insertions for the genomic-rearrangements, broad host-range editing, and dual bRNA experiments. The is110-insertion-mapping repository contains scripts used to map insertions for microbial community editing. The BridgeEvaluator repository contains scripts used to design and evaluate bRNA target sites. Representative pEdit vectors will be deposited on Addgene and available at https://www.addgene.org/Brady_Cress/.

