## Supplementary figures for "Bridge recombinase enables versatile rewriting of bacterial genomes"

**The PDF file includes:**

Materials and Methods
Supplementary Text
Figs. S1 to S22
Tables S1 to S9

**Other Supplementary Materials for this manuscript include the following:**

Data S1 to S5

### Materials and Methods

Molecular cloning of bridge recombinase vectors

Individual components of pEdit vectors (e.g. promoters, selectable markers, origins of replication) were PCR amplified from existing constructs when available, and alternatively synthesized (Twist, IDT). Components were initially cloned into storage vectors via NEB Hifi Assembly. Components were then assembled into complete pEdit vectors via Golden Gate Assembly (BbsI-HF, New England Biolabs). bRNAs were synthesized as gene fragments or as overlapping oligonucleotides (IDT) and inserted into pEdit vectors via Golden Gate Assembly (BsaI-HFv2, New England Biolabs).

BAC extraction

A set of four BACs sized at 30.4, 65.7, 78.3, and 141.8 kb were obtained from the Chlamydomonas Stock Center. We inoculated a single colony into 5 ml of LB medium containing the appropriate antibiotic and incubated at 200 rpm overnight at 30°C. We collected 3 ml of the overnight culture and resuspended the pellet in 100 µl of Suspension Solution (50 mM glucose, 25 mM Tris, 10 mM EDTA, pH 8), and then added 200 µl of freshly prepared Lysis Solution (0.2 N NaOH, 1% SDS; add 1 ml 2 N NaOH to 8.5 ml H2O, mix and add 0.5 ml of 20% SDS), 150 µl of Neutralization Solution (30 ml 5 M KOAc, 5.75 ml glacial acetic acid, and add H2O to 50 ml), and incubated on ice. The cell debris was pelleted through centrifugation at 4°C. The supernatant was transferred to a new microfuge tube, 1 ml 100% EtOH was added, and the sample was centrifuged at 4°C once more. The pellet was washed with 500 µl of 70% EtOH. Liquid was carefully removed and the sample was left out to dry. The DNA was dissolved in 30 µl of water.

General *E. coli* electroporation

Plasmids and BACs were introduced into various strains of *E. coli* through electroporation. Briefly, cells were grown in LB to an OD 600 nm of 0.4-0.6. Where indicated, antibiotics and diaminopimelic acid (DAP) were added to promote growth and selection. Cells were chilled on ice, centrifuged (10min, 4000g, 4°C), washed 2-3 times in ice-cold 10% glycerol, and resuspended in 1/100 volume 10% glycerol. 50 µl of competent cells were mixed with ~100 ng of DNA and transferred to a 1 mm electrocuvette. The cells were electroporated at 1800V and recovered at 37°C for 30 m in 500 µl SOC, supplemented with DAP where indicated. Recoveries were spread on an appropriate LB agar plate (with antibiotics and DAP supplemented where indicated).

General colony PCR (cPCR) procedure

To genotype edits through PCR, individual colonies were collected into 20 µl sterile water. 10 µl of this suspension was mixed with 10 µl 25 mM NaOH, boiled at 100°C for 10 m, and diluted with 80 µl sterile water. 1 µl of this lysate was used as a template for a GoTaq (Promega) PCR reaction, according to manufacturer's protocol. Annealing temperature was maintained at 60°C and extension time at 30 sec/kb, unless otherwise indicated. PCR products were visualized on 1% agarose gels strained with SybrSafe (Thermo Fisher)

pBAC construction

A 14 kb miniBAC positive control was assembled using NEB HiFi Assembly, according to manufacturer's protocol, from three PCR products of the BAC backbone. To introduce the C-terminal half *kanR* marker into pBAC, a *Vch*CAST vector was constructed using Gibson assembly encoding *Vch*CAST (*Vibrio cholerae* CRISPR-associated transposase) components with cargo consisting of the C-terminal *kanR* marker, IS621 donor sequence, and a spectinomycin resistance marker. The verified *Vch*CAST vector was electroporated into *E. coli* MFDpir and delivered to pBAC strains via conjugation following standard protocols. Individual colonies were picked, analyzed by cPCR and Sanger sequencing (UC Berkeley Barker DNA Sequencing Core) for the T-RL junction, and patch plated to confirm simple insert products.

*Vch*CAST-edited pBACs were then extracted by custom alkaline lysis and electroporated into the electrocompetent split kanamycin *E. coli* strains (see "Split kanamycin assay strain engineering") according to "BAC electroporation." Strains were recovered and plated on LB agar supplemented with 34 µg/ml chloramphenicol, 100 µg/ml spectinomycin, and 12 µg/ml tetracycline and outgrown for 16 h at 30°C. Individual colonies were outgrown using the appropriate antibiotics, processed via alkaline lysis to extract the edited pBAC, and sent for Big Plasmid Sequencing (Plasmidsaurus) to verify pBAC sizes. Correct clones for each pBAC size were plated on LB agar supplemented with 25 µg/ml kanamycin to confirm no spurious recombination of the *kanR* half sites in the absence of IS621. Correct clones were then glycerol stocked for all pBAC sizes in the *E. coli* BW25113 background.

<u>Split kanamycin strain engineering</u>

First, a replicative vector was constructed encoding the *Sh*CAST (*Scytonema hofmanni* CRISPR-associated transposase) system with cargo consisting of the N-terminal *kanR* marker, IS621 target sequence, and a tetracycline resistance marker. The *Sh*CAST editing vector was under the control of tsSC101, a temperature-sensitive origin of replication that allows persistence at 30°C or room temperature and is cured by outgrowth at 42°C. The resulting assembly was purified with the DNA Clean and Concentrator-5 kit (Zymo Research), electroporated into electrocompetent *E. coli* MFDpir, and individual colonies sequence-confirmed using QIAprep Spin Miniprep Kit and whole-plasmid sequencing (Plasmidsaurus). Following sequence verification, the vector was electroporated into electrocompetent *E. coli* MG1655, BW25113, and NEB10beta, recovered at 30°C, and outgrown on LB media supplemented with µg/ml carbenicillin and 12 µg/ml tetracycline for 16 h at 30°C. Colonies from each *E. coli* strain were picked, outgrown at room temperature, and heat-cured at 42°C by passaging in liquid and solid media. Individual colonies were patch plated and analyzed by colony PCR (cPCR) and Sanger sequencing (UC Berkeley Barker DNA Sequencing Core) to confirm the correct simple insert. Correct clones were glycerol stocked.

<u>Split kanamycin assay conjugations, measurement of pBAC integration, and mapping of on- and off-target integrations</u>

*E. coli* MFD*pir* (Mu-free donor strain, *pir*+) cells (*54*) were electroporated with a pEdit plasmid encoding an aTc-inducible IS621 recombinase. To measure on-target recombination frequency, we used kanamycin resistance (as conferred by the reconstituted split kanamycin marker) as our readout. The split *kanR* recombination junction was encoded in a permissive loop of the aph(3')Ia enzyme. The recipient strain encodes the target sequence at the conserved genomic safe site *caiF*. Inducing expression of pEdit in a pBAC-carrying recipient strain inserts the full pBAC into the genome, reconstituting the *kanR* marker (provided the insertion is on-target).

In all split kanamycin experiments, the pEdit plasmid was delivered by conjugation from the MFD*pir* donor strain to the recipient strain carrying pBAC. Recipient cells were cultured in biological triplicate for 16 h with shaking at 200 rpm in LB media supplemented with 12 µg/ml tetracycline, 100 µg/ml spectinomycin, and 34 µg/ml chloramphenicol at 30°C to ensure faithful replication of pBAC. Donor cells were cultured in biological triplicate per bRNA for 16 h with shaking at 200 rpm in LB media supplemented with µg/ml carbenicillin and DAP at 30°C. Cultures were mixed as previously described, spotted onto LB agar supplemented with 2 nM aTc inducer and DAP, and allowed to conjugate for 4 h at 37°C. Conjugation spots were then scraped and 100 µl resuspension was inoculated in 5 ml of LB media supplemented with chloramphenicol and carbenicillin to maintain pEdit and pBAC. Cultures were then passaged with shaking at 200 rpm for 2 h at 37°C to recover. Next, 200 nM aTc was added to induce bridge recombinase expression off pEdit and cultures were induced and passaged for 20 h at 200 rpm and 37°C. 200 µl of the passaged and induced cultures were aliquoted, serially diluted, and plated on LB agar supplemented with 25 µg/ml kanamycin (to select for recombinants) and LB agar supplemented with chloramphenicol and carbenicillin (to select for pEdit transformants). Individual colonies were analyzed by cPCR and PCR amplicon sequencing (Plasmidsaurus) to confirm the reconstituted *kanR* junction. Representative colony picks for each pBAC size were gDNA extracted using the BioSearch MasterPure™ Complete DNA and RNA Purification Kit. Purified gDNA was sent for whole-genome sequencing to confirm full pBAC integration (Plasmidsaurus).

To quantify off-target integrations, an aliquot of each conjugation mixture was seeded on LB agarose supplemented with carbenicillin and chloramphenicol and outgrown overnight at 37°C. 500 to 1,000 colonies per plate were scraped, resuspended in 1X PBS, and processed for genomic DNA extraction. Purified genomic DNA was sent for whole-genome sequencing (Plasmidsaurus). Raw fastqs from each sequencing run were used to map on- versus off-target insertions (see “Mapping of bridge recombinase insertion sites”).

<u>Targeted genomic rearrangement conjugations and measurement of inversions and excisions</u>

*E. coli* DATC (DAP-Auxotroph Transformation Conjugation, *pir+*) cells (*55*) were electroporated with a replicative RSF1010 pEdit vector encoding the IS621 recombinase, its cognate bRNA, and a hygromycin resistance cassette. pEdit donor strains were cultured in biological triplicate in LB media supplemented with 100 µg/ml carbenicillin, 50 µg/ml hygromycin, and DAP for 16 h with shaking at 200 rpm and 37°C. *E. coli* MG1655 cells were cultured in biological triplicate in LB media for 16 h with shaking at 200 rpm and 37°C. Cultures were then washed and mixed according to conjugation protocols, spotted onto LB agar supplemented with 2 nM aTc inducer and DAP, and allowed to conjugate for 4 h at 37°C. Conjugation spots were then scraped and 100 µl resuspension was inoculated in 5 ml of LB media supplemented with carbenicillin. Cultures were then passaged with shaking at 200 rpm for 2 h at 37°C to recover. Next, 200 nM aTc was added to induce bridge recombinase expression off pEdit. Cultures were induced and passaged for 16 h at 200 rpm and 37°C. 1 ml of the passaged and induced cultures were resuspended in 1X PBS, spun down, and washed in 1X PBS to remove residual rich media components. 200 µl of washed cultures were aliquoted, serially diluted in 1X PBS, and spotted on M9 agar supplemented with 200 mg/mL 2-DOG, 25 µg/ml kanamycin, 50 µg/ml hygromycin, and 100 µg/ml carbenicillin (to select for *galK* recombinants) or M9 agar supplemented with kanamycin, hygromycin, and carbenicillin (to select for transformants) and outgrown at 37°C for two days.

Individual colonies were picked and analyzed by cPCR with oligos designed to detect inversion (two amplicons primed off the inverted junctions) or excision (one amplicon primed off the new excision junction). Amplicons were sequenced using PCR amplicon sequencing (Plasmidsaurus) to validate the expected inversion and excision junctions.

Flanking colibactin biosynthetic gene cluster with *hsvTK*

An *hsvTK-specR* cassette was cloned between the terminal ends of a non-replicating *Vch*CAST expression vector. A guide RNA targeting the left end of the colibactin BGC (at *clbO)* was additionally cloned onto the vector. This vector, transformed into donor *E. coli* DATC, was introduced to recipient *E. coli* Nissle 1917 via conjugation. Donor and recipient strains were washed, concentrated 1/10, and spotted onto LB agar plates supplemented with 300 µM DAP and 200 nM aTc. Conjugation plates were incubated for 20 h at 30°C. Spots were scraped into 500 µl LB and spread on LB agar supplemented with 100 µg/ml spectinomycin plates to select for integrants. Integration was validated through PCR.

Genome-wide rearrangement assay and sequencing of recombination sites

To construct the *hsvTK* genomewide insertion library, a cassette encoding *hsvTK* and *specR* was cloned between mariner terminal ends on a non-replicating himar1C8 transposon vector. This vector was introduced to recipient *E. coli* MG1655 via conjugation. Donor and recipient strains were washed, concentrated 1/10, and spotted onto LB agar plates supplemented with 300 M DAP and 200 nM aTc. Conjugation plates were cultured for 20 h at 37°C. Spots were scraped into 5 ml LB and a fraction of this suspension spread on LB agar supplemented with 100 µg/ml spectinomycin plates to select for transconjugants. A library of 3.2e5 transconjugant CFUs was collected. gDNA was extracted from this library and AP-PCR performed (see below) to map insertion sites.

100 µl of the recipient *hsvTK* insertion library was inoculated into 1 ml LB and spectinomycin and cultured at 37°C shaking for 1 h. Bridge recombinase plasmid -containing donor strains (bRNA programmed to recombine between *galK* and *hsvTK*) were grown in LB supplemented with 100 µg/ml carbenicillin and 300 M DAP overnight at 37°C. Donor and recipient cultures were washed twice in LB and resuspended in 1/10 volume LB. 10 µl of concentrated donor and recipient were mixed and spotted onto LB agar plates supplemented with 300 M DAP and 2 nM aTc for 4 h. Spots were scraped into 5 ml LB and 100 µg/ml carbenicillin and 200 nM aTc and cultured at 37°C shaking for 20 h to induce recombination. The overnight culture was washed twice in PBS. 10X serial dilutions were performed and 5 µl of dilution series spotted onto M9 minimal media agar plates, supplemented with 0.4% glycerol and 0.25 mg/ml Thiamine-HCl, and selected with 2 mg/ml 2-Deoxygalactose (2-DOG), 100 nM 6-(β-d-2-deoxyribofuranosyl)-3,4-dihydro-8H-pyrimido [4,5-c][1,2] oxazin-7-one (dP), and 100 µg/ml carbenicillin. In parallel, washed cultures were concentrated and spread with glass beads onto the selective M9 agar plates.

96 individual colonies from the selective M9 plates were picked and screened via AP-PCR to map recombination sites, of which 67 clones had unambiguous mappings. 13 clones were further validated through Nanopore-based whole genome sequencing (Plasmidsaurus). A programmed inversion on a circular genome is defined by two arcs (major and minor) summing to the full chromosome length; the maximum effective inversion is therefore half the genome, where the two arcs are of equal length. We achieved programmed inversion of 2.35 Mb, within ~1% of the theoretical maximum effective inversion.

IS621 delivery and editing in bacterial isolates

Bacterial strains were cultured under conditions specified in Table SBB. Anaerobic species were grown in an Anaerobe Systems AS-500 chamber (5% $CO_2$, 5% $H_2$, 90% $N_2$). Editing plasmids (pEdit) carrying appropriate promoter and selection markers were customized for each phylum (**Fig. 2A, Table SBB**).

Delivery of pEdit was performed by conjugation, electroporation, or natural competence, depending on species (**Table SBB**). For conjugations, *E. coli* DAP-auxotroph strains carrying pEdit plasmids were grown in LB + 300 M diaminopimelic Acid (DAP) + 100 g/ml carbenicillin, and recipient strains under conditions specified in Table SBB. For each culture, a volume corresponding to OD 2.0/ml was centrifuged (10000 g, 1 m, RT), washed twice, and resuspended in 1/10 volume in respective recipient media. Donor and recipient cells were mixed at equal volumes (10 µl), spotted onto recipient agar media with 30 M DAP and 200 nM anhydrotetracycline (aTc) for IS621 induction. For *aerobic* strains, conjugations were incubated at 30°C for 24 h. For *anaerobic* strains, all washes were conducted anaerobically, and conjugations incubated at 37°C for 24 h in the anaerobic chamber. Conjugation spots were then collected into 500 µl of respective recipient media for downstream analysis.

For electroporations and natural competence, pEdit plasmids were prepared from *E. coli* DATC via midiprep (Zymo). For *C. glutamicum (56)*, an overnight culture was grown in BHI at 30°C. The next day, the culture was diluted to an OD600 of 0.3 into NCM medium (17.4 g/l $K_2HPO_4$, 11.6 g/l NaCl, 5 g/l Glucose, 5 g/l Tryptone, 1 g/l Yeast extract, 0.3 g/l Trisodium citrate, 0.05 g/l $MgSO_4$ * $7H_2O$, 91.1 g/l Sorbitol, 36 g/l Glycine, 4.6 g/l Isonicotinic acid hydrazine, 0.11% (v/v) Tween 80, 1.11% (w/v) DL - Threonine, pH 7.2). Cultures were incubated, shaking at 30°C, until an OD of 0.5–1.0 was reached. Cells were chilled on ice, harvested, and washed 3–4 times with ice-cold 10% glycerol by centrifugation (4,000 g, 10 m, 4°C). Cells were finally resuspended in 10% glycerol to a final OD of 20. 500 ng of plasmid DNA was added to 80 µl of electrocompetent cells and transferred to a pre-chilled 1 mm electroporation cuvette and electroporated (1.8 kV, 10 µF, 600 Ω) using a BioRad Gene Pulser Xcell. Immediately following electroporation, 920 µl of preheated BHI + 91 g/l Sorbitol + 200 nM aTc was added and transferred to a culture tube. The mixture was incubated at 46°C for 5 m, then incubated at 30°C for 2 h with shaking at 200 rpm.

For *B. subtilis* and *E. faecalis (57)*, an overnight culture of each recipient was grown in LB at 37°C. The next day, 2.6 ml the culture was diluted into 40 ml LBSP media (85 ml LB + 5 ml 10 M Sorbitol + 5 ml 1 M KH2PO4 + 5 ml 1 M K2HPO4) and cultured shaking at 37°C. At OD 0.5, 5 ml of 10x cell wall weakening solution (6.4% Glycine, 10.2% dl-threonine, 0.5% Tween 80) was added and cultured further for 1 h. Cultures were chilled on ice and harvested through centrifugation (4000 g, 10 m, 4°C). Cells were resuspended in 1 ml electroporation buffer (0.5 M trehalose, 0.5 M sorbitol, 0.5 M mannitol). Cells were washed 4x in 1 ml electroporation buffer (10000 g, 1 m, 4°C) and resuspended in 1 ml electroporation buffer. For each transformation, 60 µl electrocompetent cells were mixed with 250 ng plasmid, and transferred to a prechilled electroporation cuvette (0.1 cm gap) and electroporated (2000 V with 5.1 ms fixed time constant). Cells were immediately recovered in 4 ml LBMS (LB + 0.5 M mannitol, and 0.38 M sorbitol) + 100 µM IPTG overnight shaking at 37°C.

For *D. radiodurans (58)*, an overnight culture was grown in TGY at 30°C. The next day, the culture was diluted to an OD600 of 0.2 into fresh TGY medium and cultured, shaking, until OD 1.0. 6.66 ml culture was harvested, and 300 µl 1 M CaCl2 + 3.33 ml ice-cold 30% glycerol was added. 600 µl of this mixture was transferred to an eppie tube for each transformation. 1 µg plasmid + 6 µl 10 mM IPTG was added, and incubated on ice for 1 h, then shaking at 30°C for 1 h. 3 ml of TGY was added and cultures incubated overnight at 30°C to recover.

Editing efficiencies were quantified by plating 10x serial dilutions of post-transformation suspensions onto respective recipient agar plates with and without appropriate antibiotic (Table SBB). To increase the limit of detection, 50 µl of suspension was directly plated onto antibiotic plates. Plates were incubated under appropriate conditions until colonies were visible. For *B. subtilis* and *D. radiodurans*, individual transformants were subjected to a second round of IPTG induction (see **Fig. S7**).

Editing outcomes were assessed by colony-level and population-level analyses. Individual transformant colonies (n=11 per condition) and a wild-type control were screened via PCR for presence of pEdit and insertion at the *16S rRNA* locus (**Data S1)**. For each condition, a representative 16S-edited clone was validated by WGS with Oxford Nanopore (Plasmidsaurus). For population-level analysis, of >200 transformant colonies from selective agar plates were pooled, gDNA extracted (Biosearch Technologies Masterpure kit), and insertion sites amplified by AP-PCR for sequencing-based mapping (see below).

<u>Mapping of insertion sites through AP-PCR</u>

To identify and map insertion sites, metagenomic DNA samples were subjected to arbitrarily-primed PCR (AP-PCR) assay using an insert-specific primer and degenerate-sequence primers as described before (citation). Briefly, for genome-wide, host range, and dual bRNA editing, Q5 HotStart PCR Master Mix (NEB) was used; and for metagenomic editing experiments, GoTaq PCR Master Mix (Promega). The thermocycling conditions of the first amplification round were: 95°C for 120 s; 6 cycles of 95°C for 30 s, 30°C for 30 s, and 72°C for 60 s; 30 cycles of 95°C for 30 s, 45°C for 30 s, and 72°C for 120 s; and 72°C for 5 min. AP-PCR products were used for the second round of amplification with the following thermocycling parameters: 95°C for 120 s; 30 cycles of 95°C for 30 s, 54°C for 30 s, and 72°C for 120; and 72°C for 5 min. AP-PCR products were sequenced on an Oxford Nanopore Technologies PromethION platform (Plasmidsaurus) to get >5,000 raw reads per sample.

Raw reads were filtered to require a mean quality score of >10. The presence of 20 bp, corresponding to either the LE or RE or of the programmed insertion, was used to identify reads containing the bridge recombinase cargo. 25 bp flanking this end was selected. Flanking regions that mapped to the original donor plasmid were omitted, presumed to be donor plasmid contamination. The remaining flanking sequences were mapped to the appropriate reference genome, requiring a perfect match of the full 25 bp flanking sequence. For genomic rearrangement experiments (Fig. 1), the reference genome was *E. coli MG1655* in which *rRNA* operons were excised and consolidated into a single extrachromosomal contig to facilitate mapping to this repetitive region. For broad host-range experiments (Fig. 2), similarly, the *16S rRNA* genes from each reference genome were excised into a single extrachromosomal contig. For each mapped flanking region, the central 14 bp of the insertion site were imputed and counted. To correct for nanopore mutations, insertion sites within 5 bp were consolidated into a single insertion site assigned to the majority site.

For metagenomic editing experiments, raw reads were processed using a custom pipeline. Briefly, reads were filtered to have an average quality score > 20. Reads containing the bridge recombinase cargo were identified based on the presence of a 28 nt conserved sequence in the LE, then trimmed to remove the cargo. 16S sequences were extracted to a fasta file using barrnap. Trimmed reads were mapped to the 16S sequences using BLASTn, and all reads with no hits to the 16S sequences were mapped with Minimap2 to the provided genomes (MAGs for infant gut community and isolate WGS for adult community). For each mapped region, the central 14 bp of the insertion site were imputed and counted.

For all experiments, imputed 14 bp insertion sites were categorized by Levenshtein distance**,** (default calculation with python package RapidFuzz 3.14.5) as "target-like" or "donor-like", based on their similarity to either programmed donor(s) or target(s). Sites that were >5 Levenshtein distance from either programmed site were categorized as "unclassified". Insertion sites in each category were aligned with MAFFT using generous gap penalties (gap-open penalty = 0.05; gap-extend penalty = 0.05). Aligned sequences were used to generate a sequence logo (a 1% read frequency cut-off was used to facilitate interpretable visualization).

Development of enriched human gut communities

*Infant gut community*

A stool from a healthy infant donor (no. 1457694; UCSF–Lynch Lab; TIPS clinical trial–NCT00113659) (*59*), aged 12 months, was used as the inoculum. Briefly, 250 mg of stool was resuspended in 500 µl of sterile phosphate-buffered saline (PBS; Thermo Fisher Scientific) and homogenized by pipetting in an anaerobic chamber (90% $N_2$, 5% $CO_2$, 5% $H_2$). An aliquot of 15 µL was inoculated into 3 ml pre-reduced BHI medium in a 24-deep-well plate and incubated anaerobically at 37 °C with shaking (100 rpm) for 48 h. The culture was then passaged three times (48 h per passage) into fresh BHI medium (1:100 dilution) and incubated under the same conditions. Enriched communities were preserved as 25% glycerol stocks at −80 °C for downstream experiments.

*Adult gut community*

A small intestinal tissue specimen was collected aseptically from an adult patient with ulcerative colitis undergoing ileostomy closure. To minimize environmental contamination and oxygen exposure, the excised bowel ends were sutured immediately after resection. The sample was transferred to a biosafety cabinet and processed within 1 h. Tissue was rinsed with sterile PBS to remove residual blood, opened longitudinally, and the mucosal layer was gently scraped using sterile glass slides. Approximately 500 µl of mucosal material was transferred into an anaerobic chamber, resuspended in 1 ml sterile PBS, and homogenized by repeated pipetting. A 100 µl aliquot was inoculated into 9.9 ml pre-reduced Gifu Anaerobic Medium (GAM; HiMedia) and incubated anaerobically at 37 °C for ≥12 h. Cultures were then preserved as 30% glycerol stocks at −80 °C. For expansion, 100 µl of glycerol stock was inoculated into 1 mL modified GAM (Shimadzu 05433) and incubated anaerobically at 37 °C for 24 h. Cultures were passaged twice at a 1:10 dilution with 24 h incubation per passage. After the third passage, enriched communities were stocked as 15% glycerol stocks and stored at −80 °C for downstream editing experiments.

Conjugation and editing of microbial communities and analysis of edited cells

For conjugations, donor *E. coli* strains were cultured in LB media (BD) supplemented with DAP (60 µg/ml) and carbenicillin (carb) (100 µg/ml) in aerobic conditions at 37°C overnight. Communities were grown anaerobically at 37°C for 20 hours in Gifu anaerobic modified medium (mGAM) (Schimadzu). Before conjugation, donor *E. coli* cultures were diluted and grown to mid-log phase (OD~0.5). Then, 1 OD unit of donor cells was harvested and washed twice in anaerobic mGAM medium in the anaerobic chamber. This was added to 1 OD unit of the recipient community, and the mixture was plated on mGAM agar with DAP and 100 µM of IPTG or 200 nM of ATC for induction of bridge recombinase expression. Plates were incubated anaerobically at 37°C for ~20 h. After conjugation, growth was scrapped off the plate into 1mL of mGAM for downstream analysis. Total cell counts in the conjugation suspension were measured with a BactoBox (SBT Instruments).

For quantification of culturable and edited cells, conjugation suspensions were plated onto selective agar plates with and without antibiotics; since the plasmid encoding antibiotic resistance genes is non-replicative, genome insertion is required for cell survival in presence of antibiotics. MacConkey agar plates (MC) were supplemented with kanamycin (50 µg/ml) and chloramphenicol (34 µg/ml) and incubated aerobically at 37°C overnight. Bacteroides Bile Esculin Gentamicin agar plates (BBEG) (HiMedia) were supplemented with erythromycin (25 µg/ml) and thiamphenicol (15 µg/ml), and incubated anaerobically at 37°C for 48-72 h.

For downstream colony analysis, individual colonies were first patched-purified onto LB agar plates with kanamycin and chloramphenicol, or BBEG plates with erythromycin and thiamphenicol. For individual colony analysis, patch-purified colonies were resuspended in 20 µl of 10 mM NaOH, lysed by incubating at 100°C for 10 min, and resuspended in 80 µl of H2O. Cell debris was pelleted and 5 µl of lysate supernatant was used for downstream PCR analysis. For junction cPCR analysis at the *16S rRNA* locus, 10 colonies from each group were screened. For metagenomic DNA analysis, at least a thousand colonies from each selective media plate with and without antibiotics (except for BBEG with antibiotics that yielded tens of colonies) were scraped off and resuspended in 0.5 ml of PBS. Genomic DNA extractions were performed following the DNeasy PowerSoil Kit (Qiagen) with heating lysis step.

Metagenomic 16S rDNA amplicon sequencing and analysis

Metagenomic samples were PCR-amplified using universal forward and reverse primers annealing to the 16S rDNA V1 and V9 regions (oAOB003/oAOB004). Using Q5 High-Fidelity DNA Polymerase (NEB). Thermocycling conditions were: 98°C for 30 s; 30 cycles of 98°C for 10 s, 61°C for 20 s, and 72°C for 45 s; and 72°C for 2 min. Products were resolved by 1% TAE agarose gel electrophoresis and visualized by staining with SYBR Safe (Thermo Fisher Scientific). Sequencing was done on an Oxford Nanopore Technologies PromethION platform (Plasmidsaurus) to get at least 5,000 raw reads.

For each sample, raw reads were processed with the Pike full-length *16S rRNA* pipeline to generate operational taxonomic unit (OTU) consensus sequences (*60*). Taxonomy was assigned to each OTU consensus sequence by BLASTn (v2.13+) against the SILVA 138 SSU NR99 reference database (*61*). For each sample, OTU counts from three biological replicates were aggregated and converted to relative abundance. Community composition plots were generated with the ggalluvial R package.

TRADE editing with dual bRNAs

For each reaction, a non-replicating donor plasmid was constructed, expressing the bridge recombinase, *catP* cargo flanked by two distinct donor sites, and corresponding bRNAs. Donor *E. coli DATC*, carrying this plasmid, and recipient strains were inoculated into 3 ml LB, and incubated overnight at 37°C. A culture volume corresponding to OD 2.0/ml was centrifuged (10000 g, 1 m, RT), washed twice, and resuspended in 1/10 volume in LB. 10 µl donor and recipient suspensions were mixed and spotted onto LB agar supplemented with 30 M DAP + 2 nM aTc for induction. Conjugations proceeded for 4 hours and spots were then collected into 500 µl LB. These suspensions were inoculated at a 1:50 dilution into 3 ml LB + chloramphenicol and cultured at 37°C for 2 h. aTc was added to 200 nM and cultures further incubated for 18 h. To map insertion sites, 500 µl of induced cultures were collected, gDNA extracted (MasterPure™ Complete DNA and RNA Purification Kit), insertions amplified by AP-PCR, and deep sequenced (Plasmidsaurus ONT). Additionally, these cultures were serially diluted 10x in PBS, and 5 µl of the dilution series spotted onto selective (chlor + 250 nM dP) and nonselective (chlor) LB agar plates for efficiency quantifications. Plates were incubated at 37°C until visible colonies appeared. Further, 100 µl of appropriately-diluted cultures were spread with glass beads on selective (chlor + 250 nM dP) LB agar for phenotypic fluorescence scoring, or genotypic verification by colony PCR.

Swapping target core sequences in *mRFP*

To test the impact of orthogonal cores (Fig. 4E-H), the native "CT" core sequence in the *mRFP* target (GGACATCCTGTCCC) was muted at "GT" via MAGE (62). Briefly, the *E. coli* strain Qi coli was transformed with the oligo recombination plasmid pORTMAGE-1 (Addgene 72680). Transformants were cultured in LB and carbenicillin at 30°C until OD 0.5. Cultures were induced in a shaking 42°C water bath for 15 m, washed 3x in ice-cold 10% glycerol, and concentrated 20x in 10% glycerol. 50 µl electrocompetent cells were mixed with 1 µl 100 µM phosphorothioated mutagenesis oligo (c*t*gaaagttaccaaaggtggtccgctgccgttcgcttgggacatcGtgtccccgcagttccagtacggttccaaagcttacgttaaacac), transferred to a 1 mm electrocuvette, electroported (1800 V), and recovered for 3 h in LB and carbenicillin at 30°C. 3 cycles of MAGE were performed before identifying mutant clones via Sanger sequencing. pORTMAGE-1 plasmid was cured through antibiotic-free passaging on agar plates.

Pathway capture with dual bRNAs

For each reaction, a replicative RSF1010 pRec vector was constructed with cognate bRNAs. pCapture vectors were constructed containing a F-ori2 origin of replication, a second shuttle origin of replication (pRK2 oriV with the *trfA* replication protein under the control of crystal violet-inducible pJExA1 promoter (*63*) for Pseudomonadota; or pGA1 oriV (*64*) for Actinomycetota), *catP* flanked by two donor sequences embedded within *hsvTK* and *sacB* respectively, *specR* (*aadA1*), and a constitutive promoter oriented towards the pathway capture site. pCapture was transformed into corresponding pathway-containing recipient strains via conjugation. For the capture reaction, donor *E. coli* DATC, carrying pEdit, and pCapture-containing recipient strains were inoculated into 3 ml LB, and incubated overnight at 37°C. A culture volume corresponding to OD 2.0/ml was centrifuged (10000 g, 1 m, RT), washed twice, and resuspended in 1/10 volume in respective recipient media. 10 µl donor and recipient suspensions were mixed and spotted onto respective recipient agar media supplemented with 30 M DAP + 2 nM aTc for induction. Conjugations proceeded for 4 h and spots were then collected into 500 µl LB. These suspensions were inoculated at a 1:50 dilution into 3 ml LB + chloramphenicol and cultured at 37°C for 2 h. aTc was added to 200 nM and cultures further incubated for 18 h. Optionally, cultures were backdiluted 1:50 into fresh cultures for a second 24 h round of induction. These cultures were serially diluted 10x in PBS, and 5 µl of the dilution series spotted onto selective (10 µg/ml chlor + 250 nM dP + 10% sucrose) and nonselective (chlor) LB agar plates for efficiency quantifications. Plates were incubated at 30°C until visible colonies appeared. Further, 100 µl of appropriately-diluted cultures were spread with glass beads on selective (10 µg/ml chlor + 250 nM dP + 10% sucrose) LB agar, and incubated at 30°C for genotypic verification by colony PCR.

Transfer of captured pathways

Pathways on pCapture were transferred via conjugation into *Corynebacterium glutamicum, Klebsiella michiganensis*, and *Pseudomonas simiae*. The latter two strains were labeled with gentamicin resistance to facilitate selection and quantification. Briefly, the *gentR* gene (*aacC1*) was delivered via *Vch*CAST into noncoding safe sites in the genome, as described above. 24 h triparental matings on LB agar at 30°C were used to conjugate pCapture from the pathway donor strain to appropriate recipients, using *E. coli MC4100* [pRK2] as the helper strain. For *K. michiganensis*, and *P. simiae*, conjugation plates were supplemented with 50 nM crystal violet (CV) to sustain the shuttle *oriV*. Transconjugants were selected on appropriate media at 30°C (*C. glutamicum*: LB agar + 50 µg/ml nalidixic acid + 600 µg/ml spectinomycin; *K. michiganensis*: LB agar + 20 µg/ml gent + 100 µg/ml spectinomycin + 50 nM CV; *P. simiae*: LB agar + 20 µg/ml gent + 100 µg/ml streptomycin + 50 nM CV).

<u>Flow cytometry analysis</u>
To phenotypically assay strains transferred with sf*GFP-mRFP-kanR*, strains were cultured at 30°C until late log growth (*C. glutamicum*: LB + 50 µg/ml nalidixic acid + 600 µg/ml spectinomycin; *K. michiganensis*: LB + 20 µg/ml gent + 100 µg/ml spectinomycin + 50 nM CV; *P. simiae*: LB + 20 µg/ml gent + 100 µg/ml streptomycin + 50 nM CV). Cultures were washed twice with PBS and diluted 1:10 into 500 µl PBS. Cells were run on a SONY SH800S cell sorter, and analyzed with the FlowCal v1.3.1 Python package. FITC filter 488 nm(Ex)/525 nm(Em) and mCherry filter 561 nm(Ex)/600 nm(Em) were used for GFP and RFP, respectively. Singlet cell populations were gated by forward and side scatter areas, and ~100 k cells were recorded per sample.

<u>Growth assays on lactose</u>
To phenotypically assay *C. glutamicum* and *K. michiganensis* transferred with *lacZY*, strains were cultured until late log growth under conditions stated above. Cultures were washed twice in PBS and diluted 1:50 into 150 µl minimal media in a 96 well plate. *C. glutamicum* was assayed in RCH2 media (*65*) (0.25 g/l $NH_4Cl$, 0.1 g/l KCl, 0.6 g/l $NaH_2PO_4$·H2O, 11 g/l PIPES, 1% (v/v) Wolfe's mineral mix ATCC MD-TMS, 1% (v/v) Wolfe's vitamin mix ATCC MD-VS, pH 7.2) supplemented with 60 mM D-glucose or 60 mM beta-lactose. *K. michiganensis* was assayed in BMC media (*66*) (6 g/l $Na_2HPO_4$, 3 g/l $KH_2PO_4$, 0.5 g/l NaCl, 1 g/l $NH_4Cl$, 0.5% (v/v) 200x Salt Mix {80 g/l $MgSO_4$·7H2O, 4 g/l $FeSO_4$·7H2O, 0.4 g/l $MnSO_4$·H20, 5 g/l NaCl}, 1% (v/v) Wolfe's mineral mix ATCC MD-TMS, 1% (v/v) Wolfe's vitamin mix ATCC MD-VS, pH 7.2) supplemented with 25 mM D-glucose or 25 mM beta-lactose. OD600 was measured kinetically on a Biotek Synergy H1 plate reader for 84 h, continuously shaking at 30°C.

### Supplementary Text

#### Expected Recombination Products with Multi-bRNA Recombination

Compared to CRISPR-based gene editors such as Cas9, Bridge Recombinase is expected to produce a quadratic number of reaction product classes when multiple guide/bRNAs are co-expressed.

Consider the expression of *n* guide/bRNAs. The gene editor can react "on-target" or "off-target". Here, we define an "on-target" reaction as one that occurs at the true sequence programmed by the guide/bRNA. We assume that an "off-target" reaction occurs at a sequence that imperfectly matches but still resembles the programmed sequence (i.e., *target-like*).

Cas9 reaction classes:
For Cas9, each gRNA can mediate an on-target and an off-target reaction class, so for n co-expressed gRNAs, we can expect:

$$ReactionClasses(n) = n\left(reaction_{on-target} + reaction_{target-like}\right) = 2n$$

Effectively, reaction class complexity scales linearly with the number of co-expressed gRNAs.

Bridge Recombinase reaction classes:
However, with Bridge Recombinase, each bRNA has two binding loops that can each recognize either a "true" sequence or its imprecise counterpart (a "-like" sequence). Furthermore, because two distinct bRNAs can occupy a synaptic complex, reactions can occur between sequences programmed on two different bRNAs. Each bRNA therefore contributes 4 recognizable sequence classes:

*target*, *target-like*, *donor*, and *donor-like*

For *n* co-expressed bRNAs, this yields 4*n* total sequence classes that can recombine in all pairwise combinations. Because two copies of the same sequence class can also recombine with each other (e.g., donor with donor), we count unordered pairs *with repetition*:

$$ReactionClasses(n) = \binom{4n+1}{2} = \frac{4n(4n+1)}{2} = 8n^2 + 2n$$

The number of possible reaction classes, therefore, scales quadratically with the number of co-expressed bRNAs.

Bridge recombinase constrained reaction classes:
However, for the specific experimental setup in **Fig. 4**, each reaction must involve one of the true donor sequences, and two true donors cannot react with each other. Each of the *n* true donors can therefore react with any of the 3*n* non-true-donor sequences (true *targets*, *target-likes*, and *donor-likes* across all *n* bRNAs):

$$ReactionClasses(n) = n\ x\ 3n = 3n^2$$

Even under the experimental constraints of **Fig. 4**, the complexity of possible reactions still scales quadratically with the number of co-expressed bRNAs, substantially surpassing the linear scaling of Cas9.

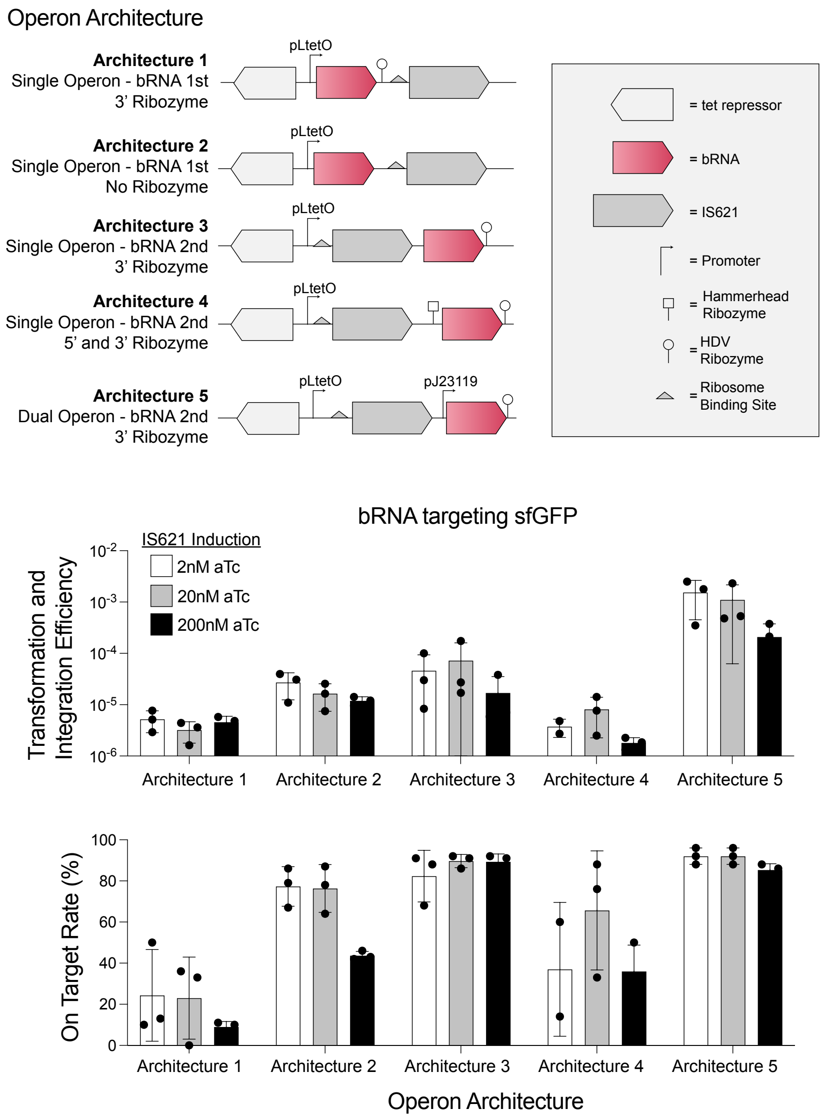


**Fig. S1. Different recombinase and bRNA expression architectures are tested, identifying dual operon architectures as most efficient for pEdit (Related to Fig. 1).** Different operon architectures of IS621 recombinase and bRNA expression were constructed and tested. The bRNA was programmed to target an insertion of *catP* into chromosomally-integrated *sfGFP* in *E. coli*. This bridge recombinase donor plasmid was conjugated into recipient cells under different induction levels IS621. Recombinants were quantified as the fraction of recipients acquiring chloramphenicol resistance. The on target rate was quantified by assaying for *sfGFP* disruption with fluorometric screening. Transformation/integration efficiencies and on target rates are plotted to visualize the impact of expression architectures and induction levels on bridge recombinase activity.

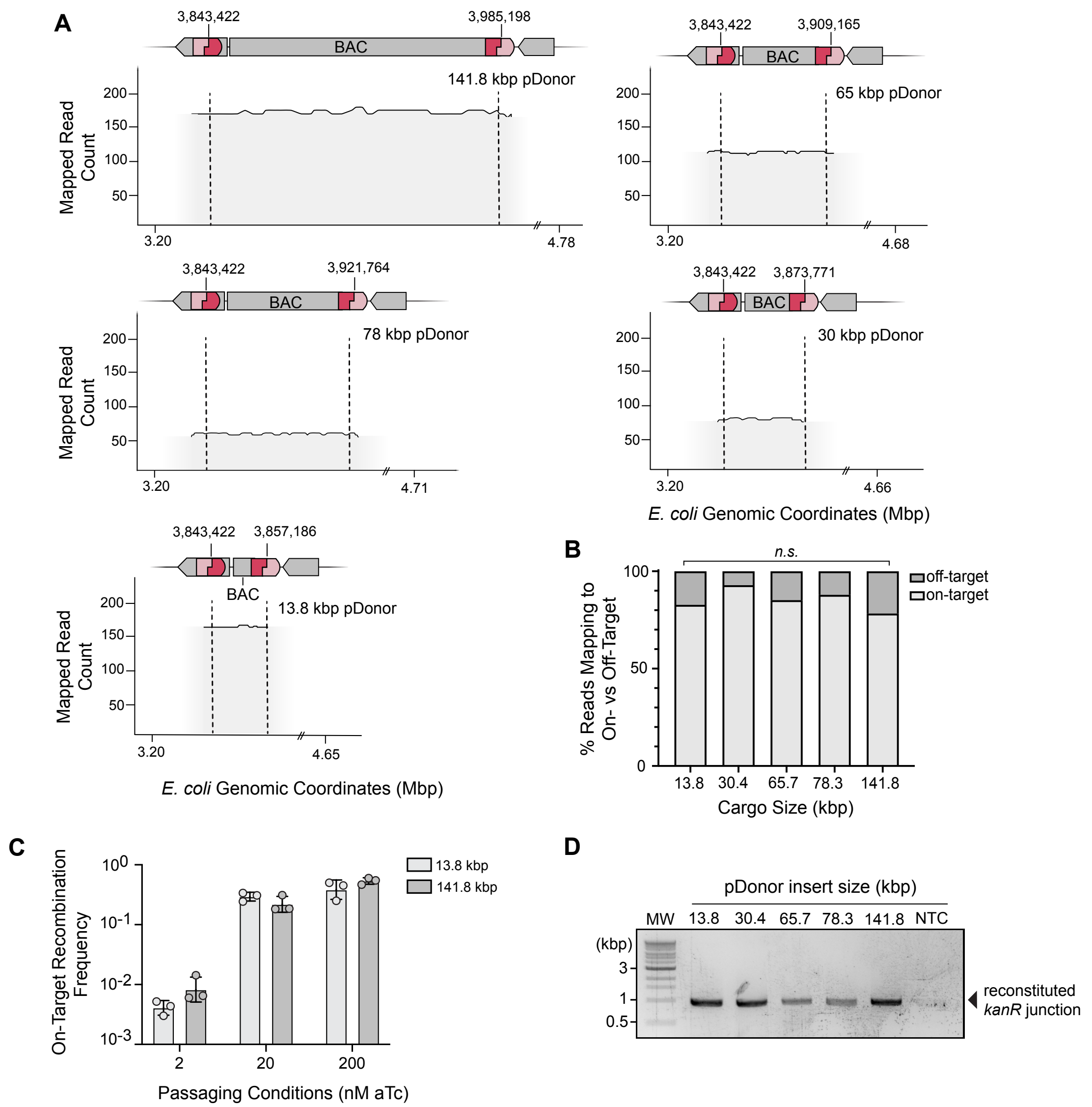


**Fig. S2. Bridge Recombinase integrates large DNA cargo (Related to Fig. 1). (A)** Representative colonies from each integrated pBAC were sent for whole-genome sequencing to validate the full on-target insert. Raw fastqs from each sequenced colony were mapped onto an *in silico* reference genome with the pBAC integrated at the IS621 target site. Mapped fastqs showed coverage of at least 75 reads across the pBAC insert at the intended target site for all pBAC vectors, demonstrating contiguous integration into the genome. **(B)** Recipient cells carrying pBAC and genomically encoding the IS621 target sequence and N-terminal *kanR* marker were passaged and induced for bridge recombinase expression. Passaged and induced cells were plated on LB agar supplemented with carbenicillin and chloramphenicol, with no selection on kanamycin-containing media for on-target recombinants. The fraction of insertions on-target and off-target are computed via deep sequencing and similarly are invariant to insertion size (one-way ANOVA, p-value > 0.9999). **(C)** During passaging, bridge recombinase expression was continuously induced with aTc. Three different concentrations of aTc were tested (2 nM, 20 nM, and 200 nM) to determine which induction level yielded the highest ratio of on-target recombinants (*kanR* CFU/ml). **(D)** cPCR of reconstituted *kanR* junction in the split kanamycin assay confirms on-target integration and expected amplicon size.

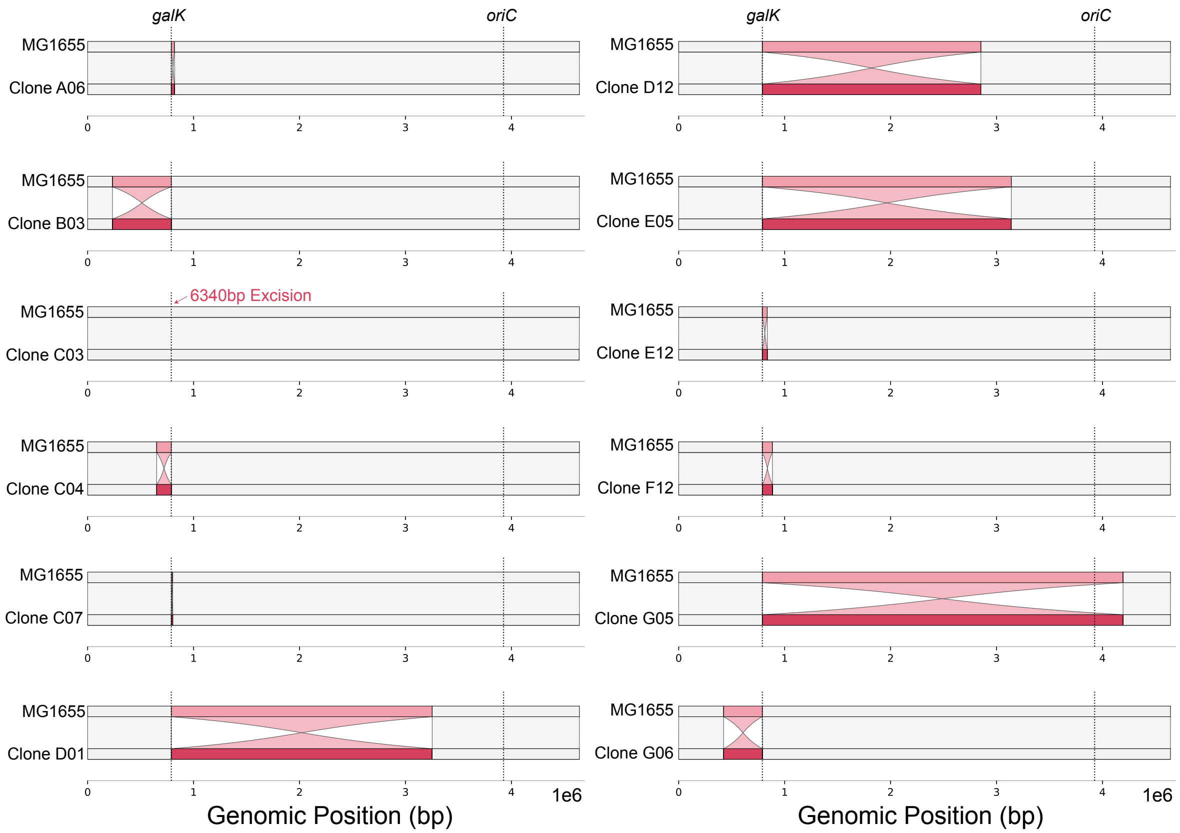


**Fig. S3. Bridge Recombinase catalyzes genome scale chromosomal rearrangements, verified by whole genome sequencing (Related to Fig. 1).** Bridge recombinase was programmed to catalyze inversions and excisions between *galK* and randomly distributed *hsvTK*. Post dP- and 2-DOG selection, 12 colonies were selected for validation through whole genome sequencing. progressiveMauve was used to align selected clones to the parental *E. coli* MG1655 genome. Observed structural rearrangements are highlighted in red. Except clone C03, all highlighted rearrangements are inversion.

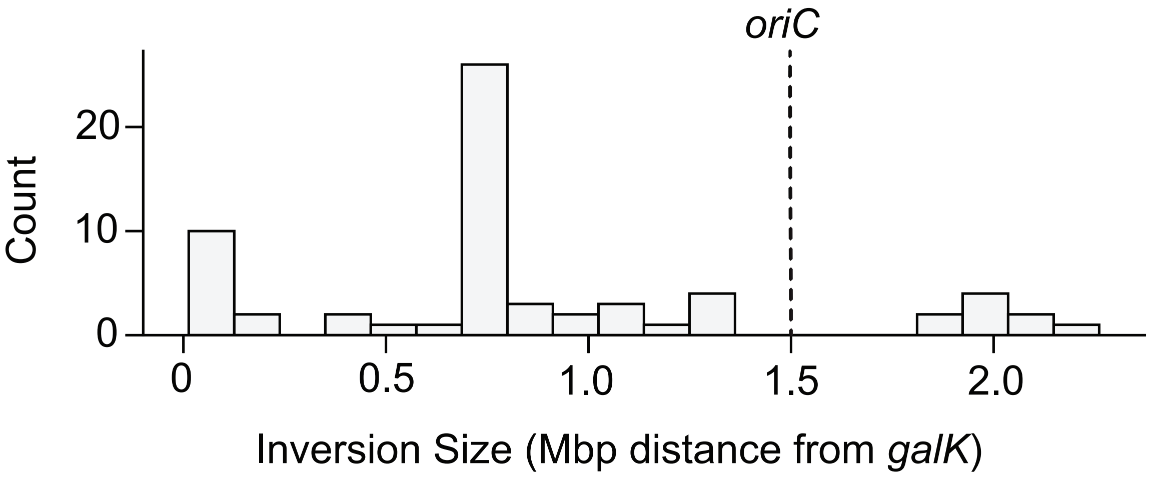


**Fig. S4. Bridge recombinase catalyzes chromosomal rearrangements with broad size distribution (Related to Fig. 1).** Bridge recombinase was programmed to catalyze inversions between *galK* and randomly distributed *hsvTK*. Post dP- and 2-DOG selection, 67 colonies were randomly screened for inversions. The frequency distribution of observed inversions are plotted as a function of inversion size.

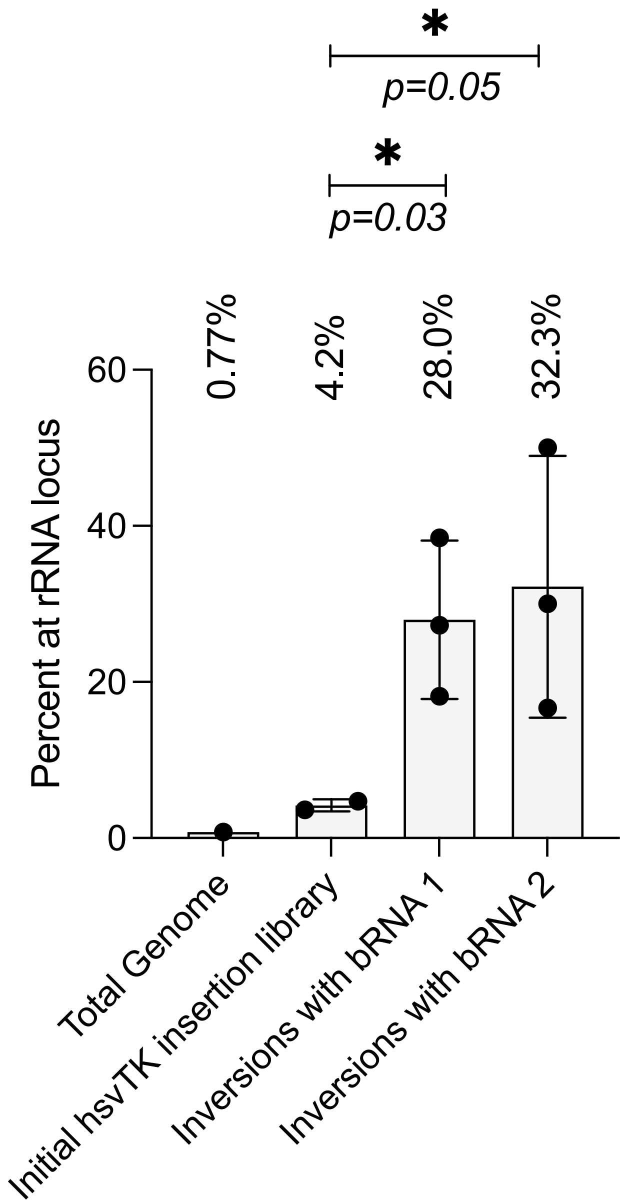


**Fig. S5. Genome-wide inversions are significantly enriched at ribosomal RNA operons. (Related to Fig. 1).** Bridge recombinase was programmed to catalyze inversions and excisions between *galK* and randomly distributed *hsvTK* in *E. coli MG1655*. Post dP and 2-DOG selection, 67 colonies were randomly picked and screened via AP-PCR to map inversion and excision sites. The distribution of *hsvTK* in the pre-recombination library was similarly mapped by AP-PCR using gDNA extracted from 500 µl of library cell culture. The fraction of inversions observed at rRNA operons was quantified, and compared to the footprint of rRNA operons within the genome, and the fraction of *hsvTK* within rRNA operons in the initial library. A one-tailed t-test is used to show bridge-mediated inversions at rRNA operons were significantly enriched compared to the initial library.

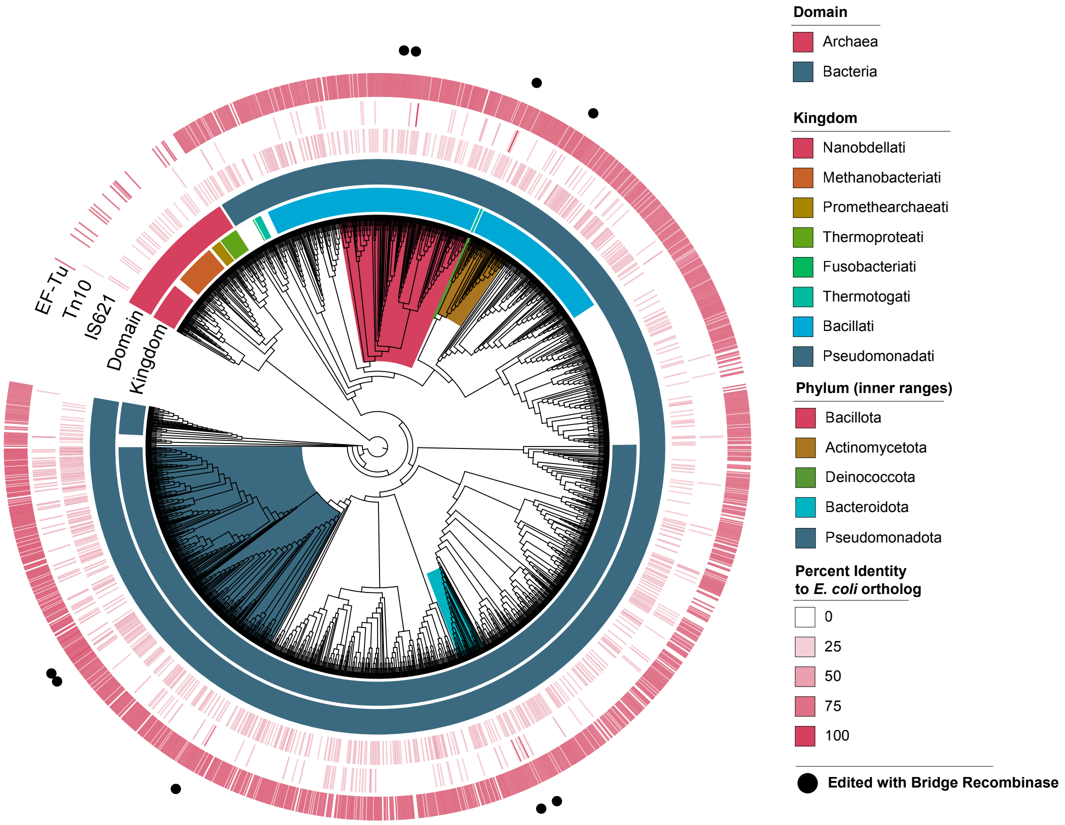


**Fig. S6. IS621 orthologs are widely distributed across the bacterial and archaeal tree of life (Related to Fig. 2).** GTDB representative bacterial and archaeal genomes (v10-RS226) were screened with mmseqs2 for homologs of the *E. coli* variant of bridge recombinase (IS621), Tn10, and EF-Tu (query coverage >0.6 and e-value < 1e-5 cutoff). The GTDB prokaryotic tree was pruned to the "order" level. Within each order, the average percent identity of retrieved homologs (vs the *E. coli* variant) are computed and plotted. IS621-like bridge recombinases are widely distributed across bacterial and archaeal taxa, compared to more narrowly constrained IS elements like Tn10.

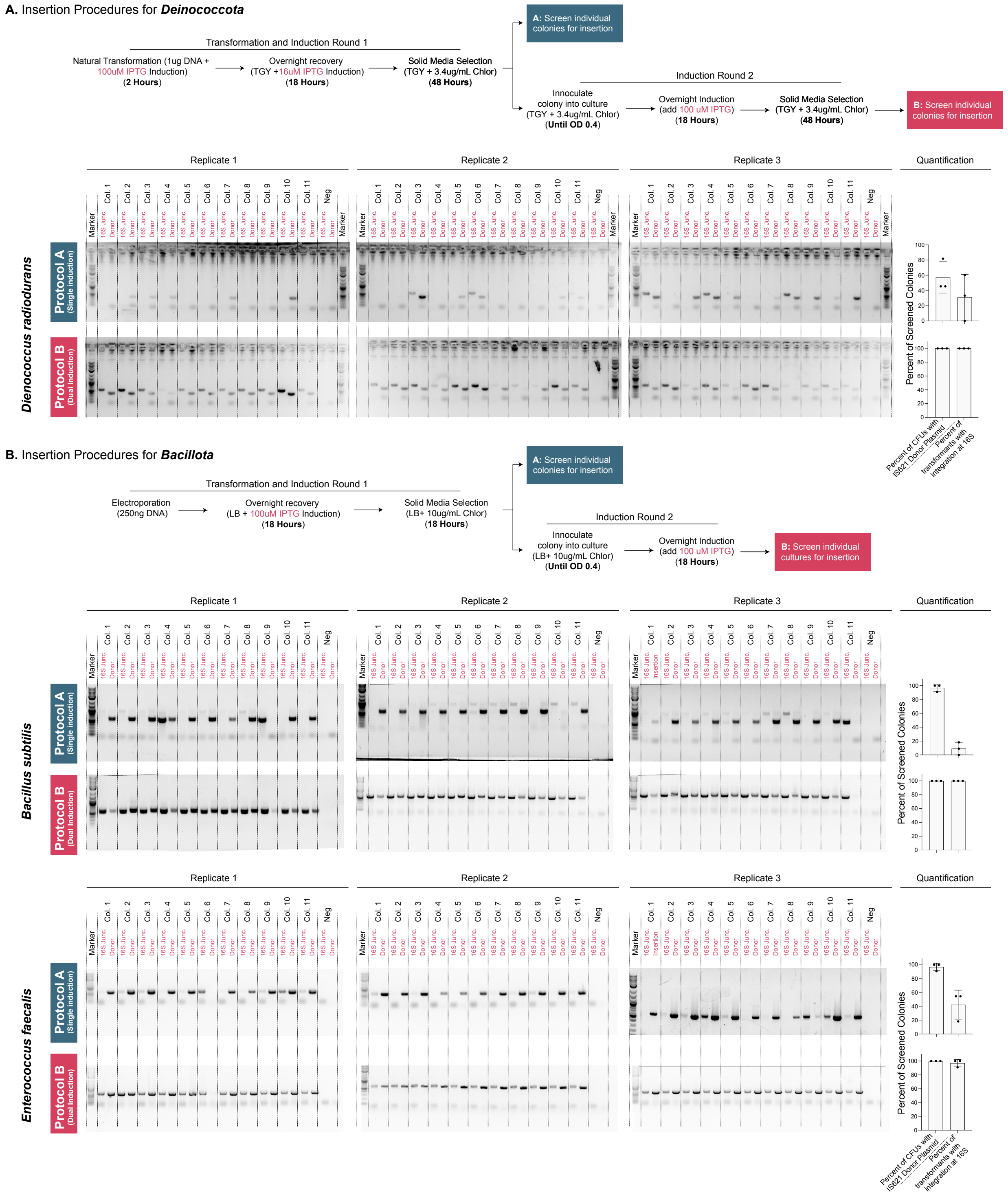


**Fig. S7. Assaying integration in *Deinococcus radiodurans*, *Bacillus subtilis*, and *Enterococcus faecalis,* which are uniquely transformed with replicating bridge recombinase vectors (Related to Fig. 2).** For each species edited with a replicating donor plasmid, two induction and selection protocols are tested. PCR is used to assess 1) the presence of donor cargo (*catP*) anywhere within the genome, and 2) the presence of integration specifically at the *16S rRNA* locus. The fraction of integrants at 16S is plotted.

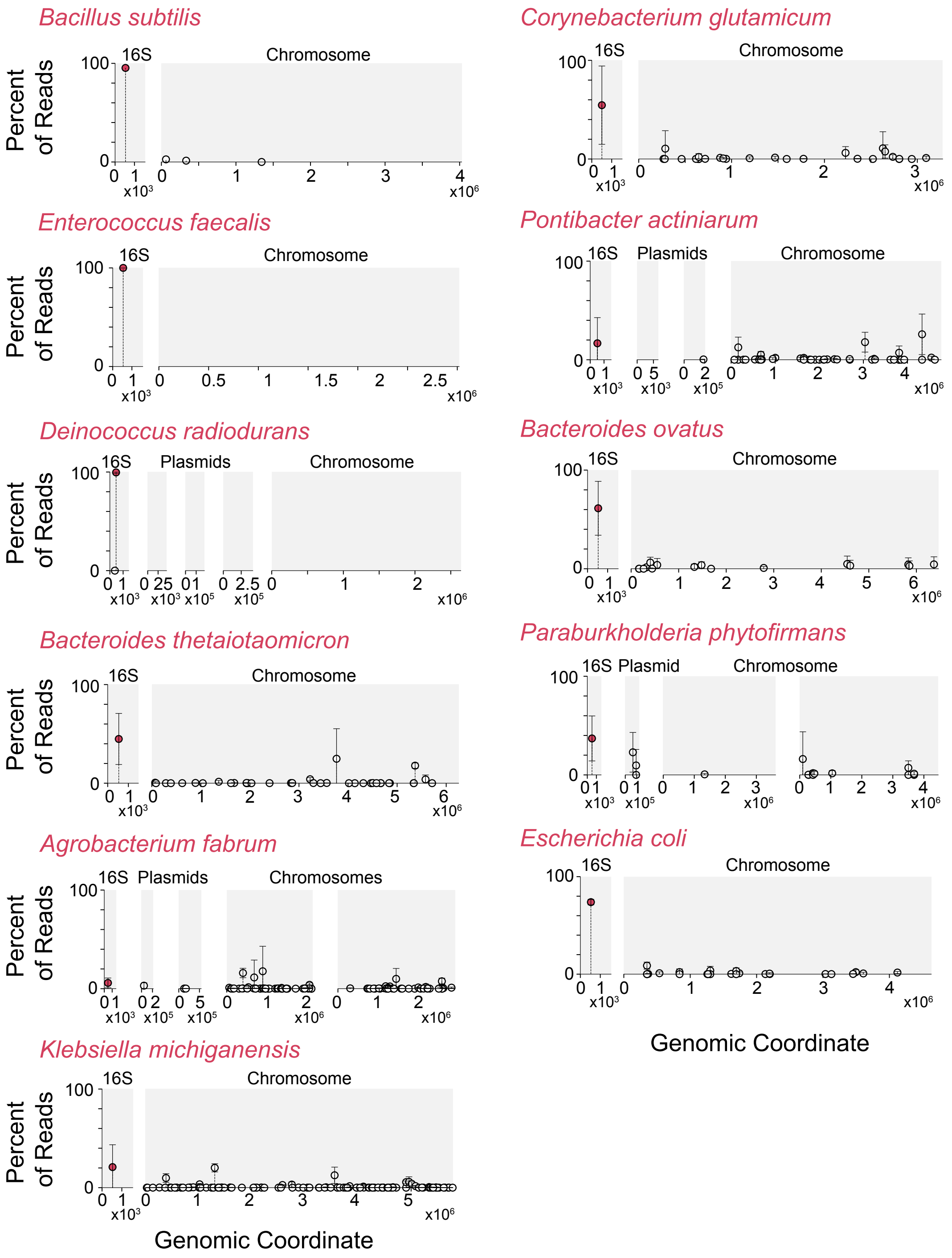


**Fig. S8. Insertion maps indicating observed integration sites across diverse bacterial species (Related to Fig. 2).** For each recipient species, bridge-mediated insertion sites are mapped with AP-PCR followed by deep sequencing. Observed insertion sites and the raw read counts are plotted. 16S rRNA genes, present in multiple copies in each tested genome, are collapsed into a single contig for mapping.

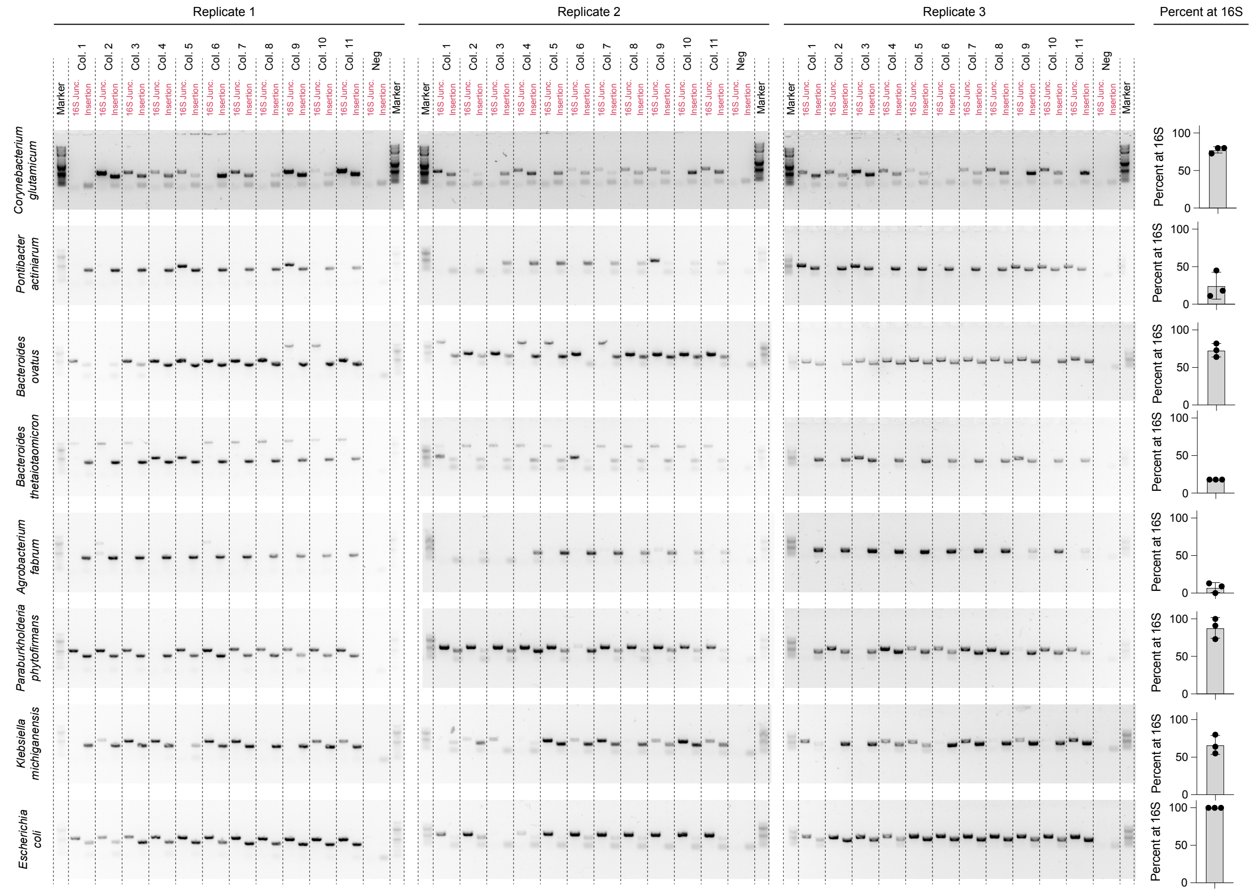


**Fig. S9. Rates of integration at the *16S rRNA* gene across edited species (Related to Fig. 2).** For each species edited with a non-replicating donor plasmid, post-integration, individual colonies are screened via PCR to assess 1) the presence of donor cargo (*catP*) anywhere within the genome, and 2) the presence integration specifically at the *16S rRNA* locus. The fraction of integrants at 16S are plotted.

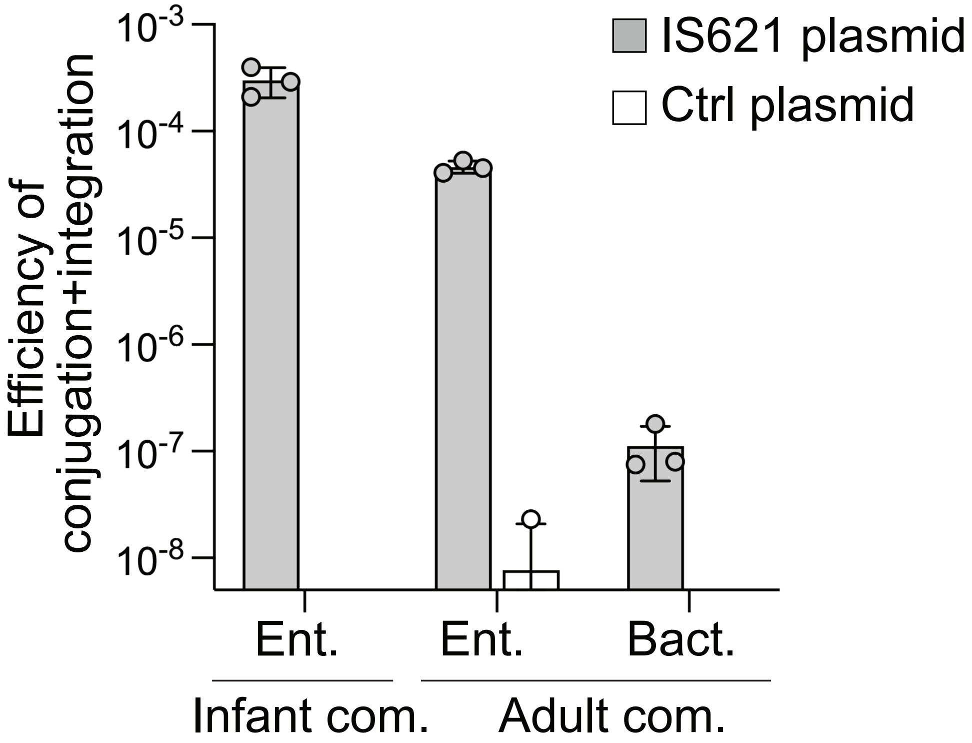


**Fig. S10. Efficiency of delivery and IS621-mediated genome integration in gut communities (related to Fig. 3).** Following delivery via conjugation of an IS621 editing plasmid or a control plasmid lacking the IS621 gene and bRNA to infant and adult gut communities, cells were plated on selective media with and without antibiotics. Because the delivery plasmid is non-replicative, antibiotic-resistant colonies require genomic integration. Transconjugant efficiency is calculated as the ratio of colony counts on selective media with antibiotics to those without. Ent., Enterobacteriaceae-selective media; Bact., Bacteroidaceae-selective media; com., community. Mean ± s.d. of three biological replicates.

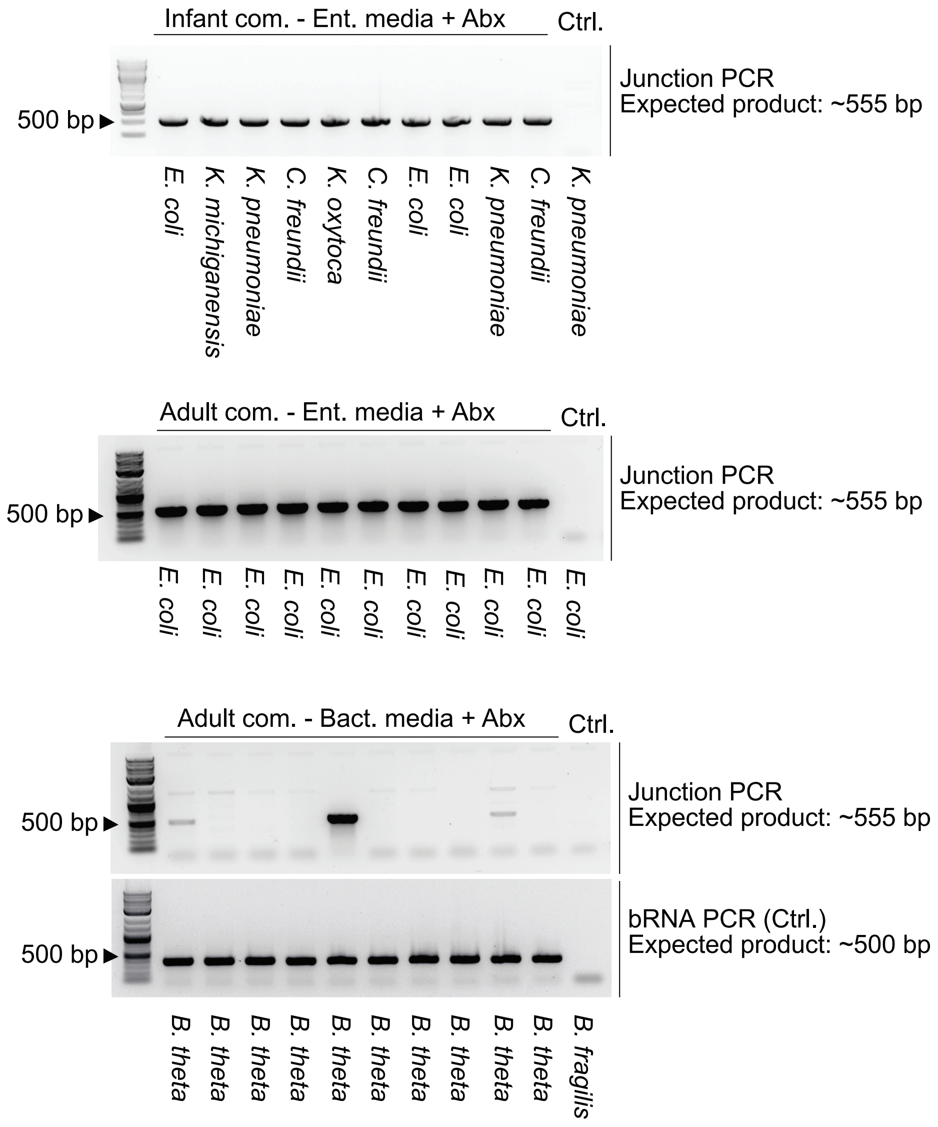


**Fig. S11. PCR confirmation of IS621-mediated integration at the 16S target site in individual community members (related to Fig. 3).** Individual colonies were picked from selective plates with antibiotics following community editing of infant (top) or adult (middle, bottom) enriched gut communities; control (Ctrl.) colonies were picked from plates without antibiotics. Junction PCR generates a ~555 bp product diagnostic of on-target integration. Species identity of each colony was determined by full-length 16S sequencing. In the Bacteroidaceae-selective media for the adult community (bottom), a bRNA PCR control (~500 bp product) confirms delivery of the editing vector. *B. theta*, *B. thetaiotaomicron*; *B. fragilis*, *Bacteroides fragilis*; Ent. media, Enterobacteriaceae-selective media; Bact. media, Bacteroidaceae-selective media; Abx, antibiotics; Ctrl., no-antibiotic control.

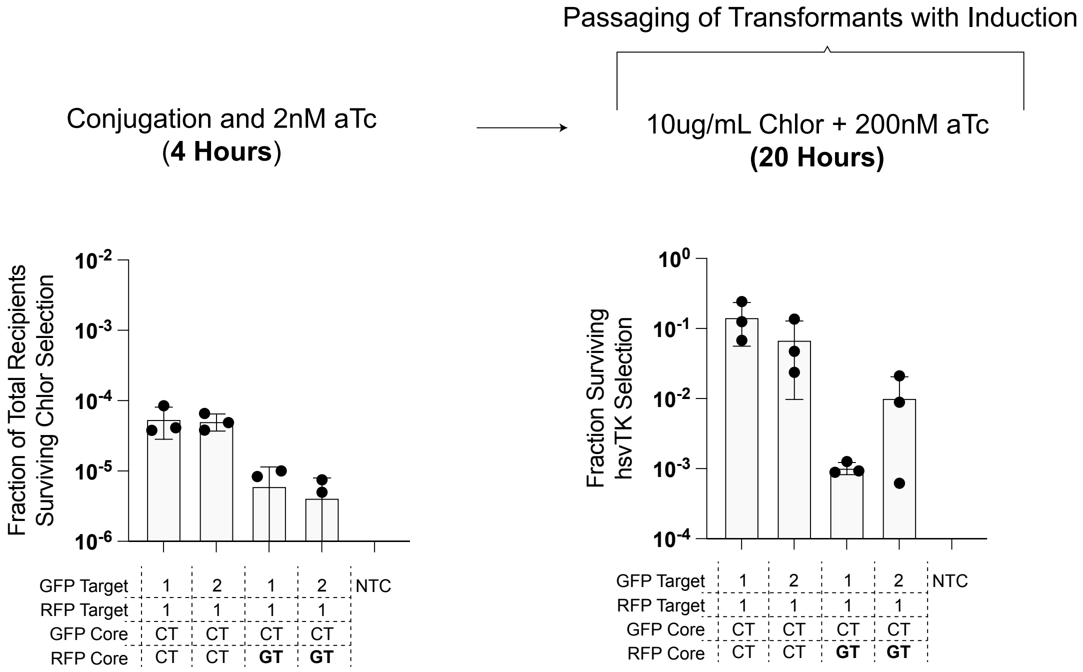


**Fig. S12. Selection methodology and raw integration and selection efficiencies for TRADE disruption of the *sfGFP-mRFP* locus with dual bRNAs (Related to Fig. 4).** For the *intragenic* disruption of GFP and RFP, transformation, induction, and selection protocols for search-and-replace editing with a non-replicating bridge donor plasmid are outlined. For each bRNA pair, fractions surviving the respective selections are plotted.

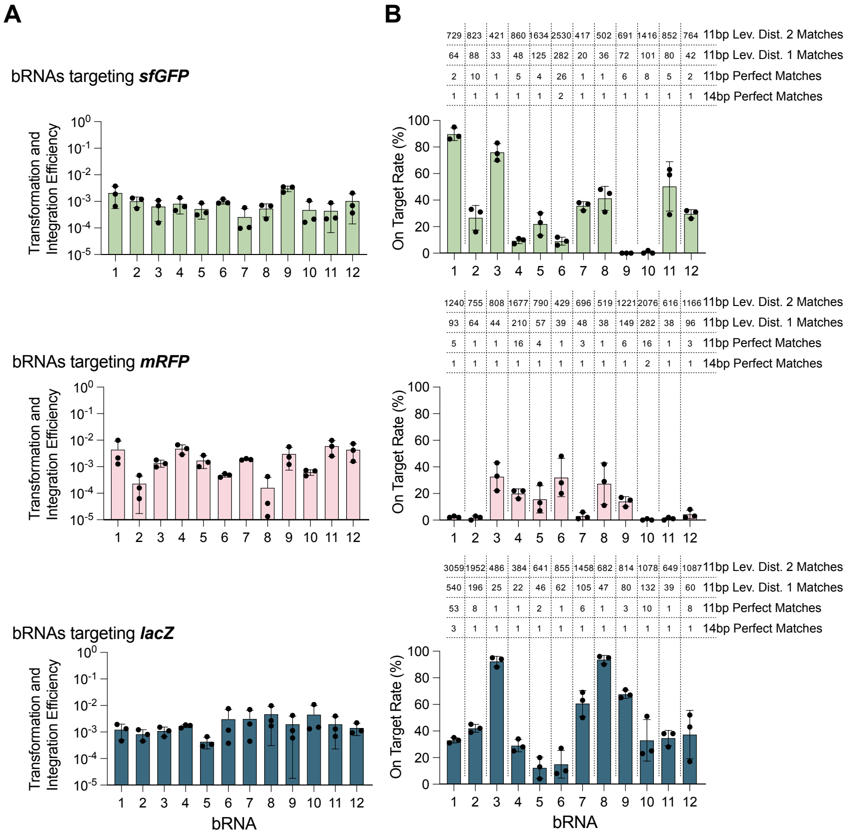


**C**

Multivariate linear regression parameters

* Regressing **"On-Target Rate"** on **"11bp Perfect Matches"**, **"11bp Lev. Dist. 1 Matches"**, **"11bp Lev. Dist. 2 Matches"**

**Pooled $R^2$:** 0.073

**Per-gene $R^2$:**

| *sfGFP* | *mGFP* | *lacZ* |
|---|---|---|
| 0.567 | 0.251 | 0.034 |

**Per-gene correlation coefficients:**

| *sfGFP* | *mGFP* | *lacZ* | |
|---|---|---|---|
| -0.47 | -0.19 | -0.18 | 11bp Perfect Matches |
| -0.38 | -0.27 | -0.18 | 11bp Lev. Dist. 1 Matches |
| -0.47 | -0.36 | -0.17 | 11bp Lev. Dist. 2 Matches |

**Fig. S13. bRNAs targeting reporter loci greatly vary in specificity, despite having similar bulk insertion efficiencies (Related to Fig. 2).** bRNAs were programmed to target an insertion of *catP* into chromosomally-integrated *sfGFP, mRNA,* or *lacZ* in *E. coli*. This bridge recombinase donor plasmid was conjugated into recipient cells under different induction levels of IS621. **(A)** Recombinants were quantified as the fraction of recipients acquiring chloramphenicol resistance. **(B)** The on-target rate was quantified by assaying for *sfGFP*/*mRFP* disruption with fluorometric screening or *lacZ* disruption with X-Gal blue/white screening. For each bRNA, the abundance of perfect and imperfect-match occurrences of the programmed target sequence in the reference *E. coli* genome is quantified (a central "CT" core sequence is required, as a fixed constraint, for each occurrence considered). **(C)** The on-target rate of each bRNA is linearly regressed on the abundance of perfect and imperfect-match occurrences of the programmed target sequence. Multivariate regressions are run for each gene and for the whole dataset. Correlation values are tabulated. The abundance of *target-like* sequences in the genome only partially explains the variability in specificity.

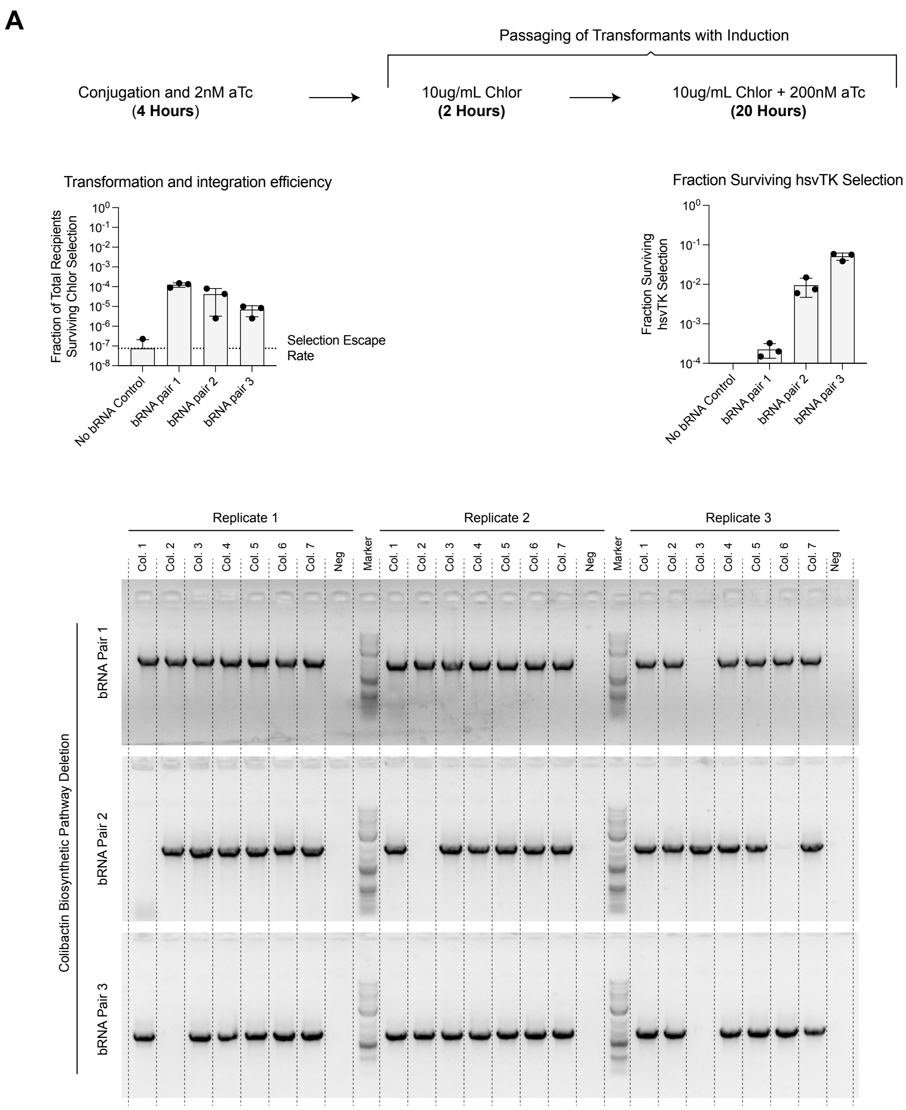


**Fig. S14. Selection methodology and raw recombination and selection efficiencies for TRADE editing of the colibactin biosynthetic locus with dual bRNAs (Related to Fig. 4).** **(A)** For the colibactin biosynthetic locus, transformation, induction, and selection protocols for TRADE editing with a non-replicating bridge donor plasmid are outlined. For each reaction, fractions surviving the respective selections are plotted. **(B)** Following search-and-replace reaction and selection, individual colonies were screened via PCR to confirm loss of the biosynthetic pathway. (expected wt amplicon = 53.5 kb; expected deletion amplicon = 1.9 kb for bRNA pair 1, 1.9 kb for bRNA pair 2, and 1.4 kb for bRNA pair 3).

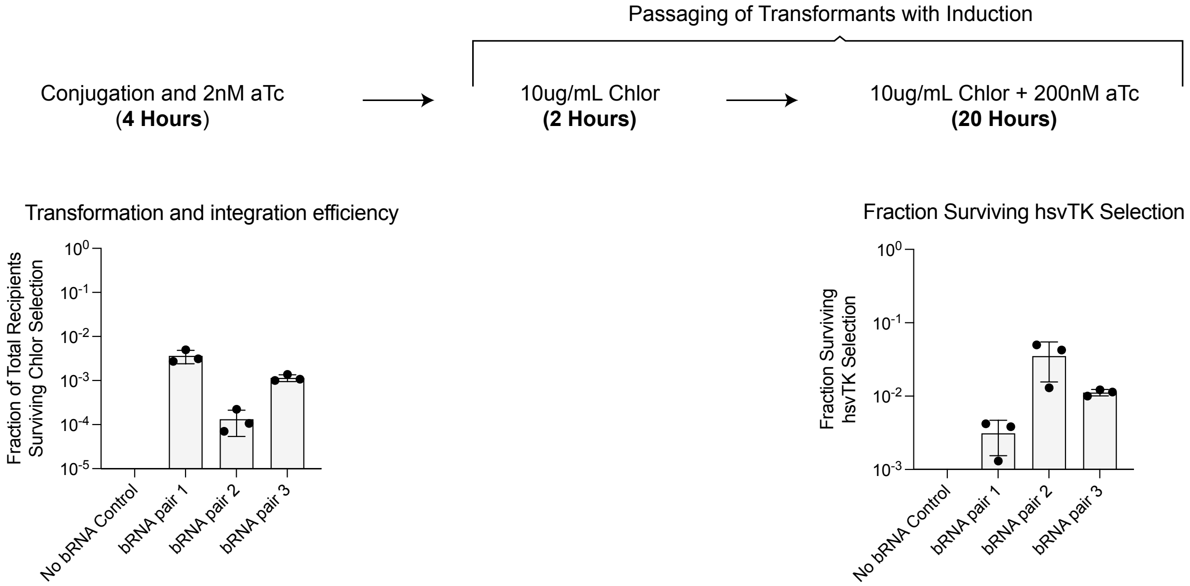


**Fig. S15. Selection methodology and raw recombination and selection efficiencies for TRADE editing of *sfGFP-mRFP-kanR* locus with dual bRNAs.** For the *extragenic* disruption of the *sfGFP-mRFP-kanR* locus, transformation, induction, and selection protocols for TRADE editing with a non-replicating bridge donor plasmid are outlined. For each bRNA pair, fractions surviving the respective selections are plotted.

**A.** Deletion of GFP RFP kanR Cluster

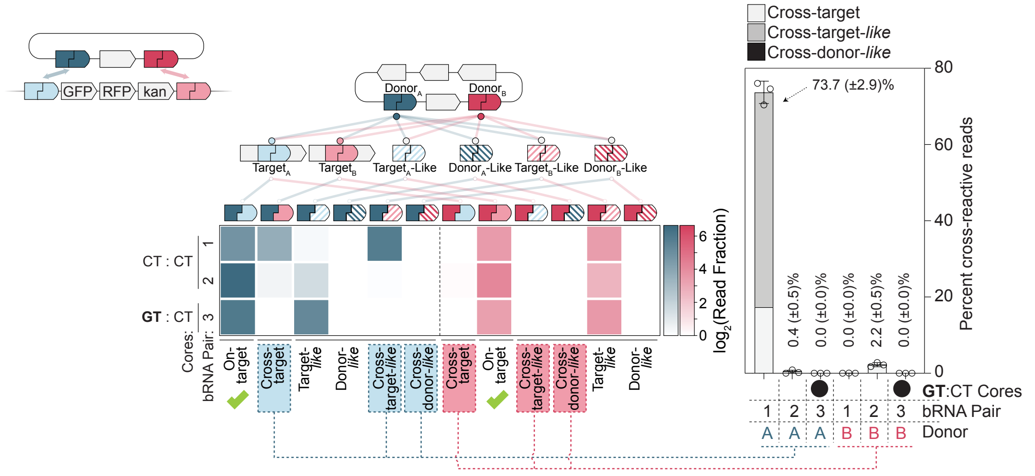


**B.** Deletion of Colibactin Biosynthetic Cluster

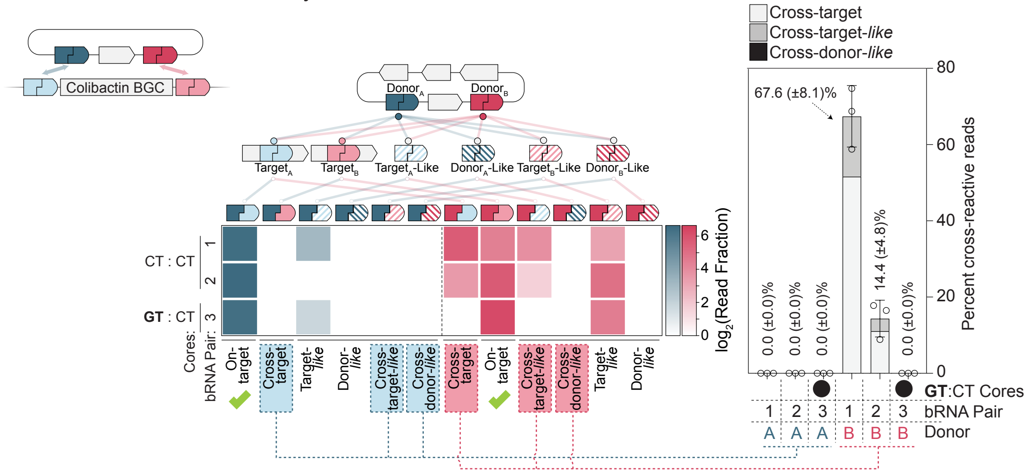

**C.**

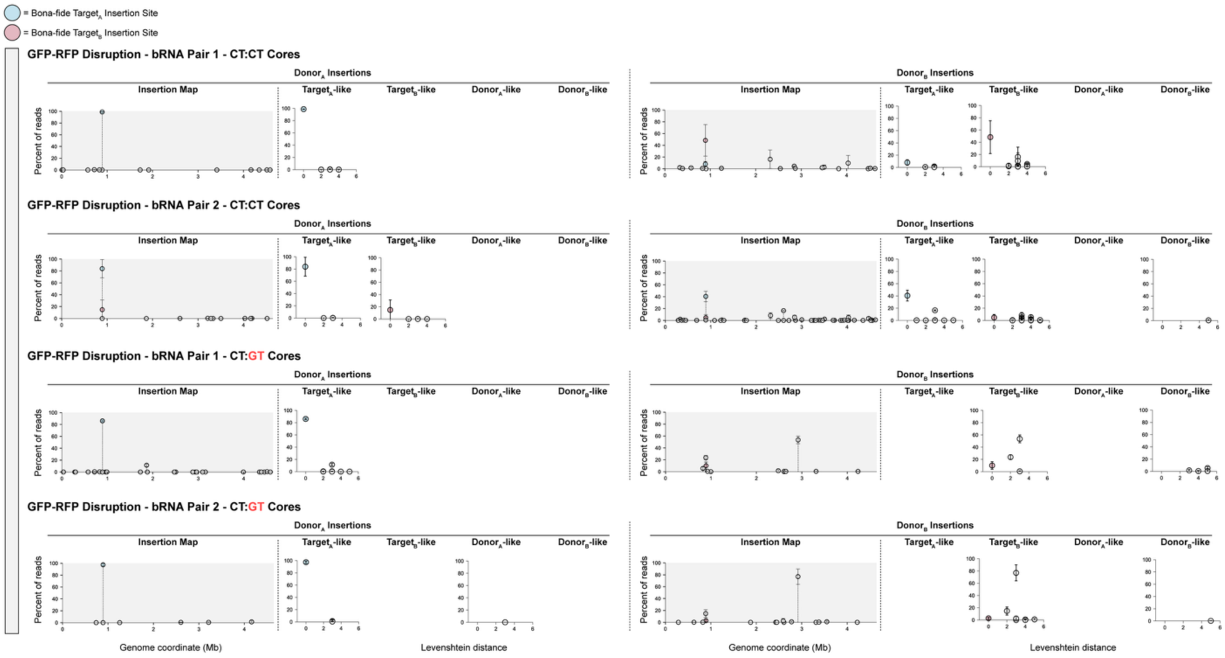
= Bona-fide Target$_A$ Insertion Site
= Bona-fide Target$_B$ Insertion Site
GFP-RFP Disruption - bRNA Pair 1 - CT:CT Cores
GFP-RFP Disruption - bRNA Pair 2 - CT:CT Cores
GFP-RFP Disruption - bRNA Pair 1 - CT:GT Cores
GFP-RFP Disruption - bRNA Pair 2 - CT:GT Cores
Donor$_A$ Insertions
Donor$_B$ Insertions
Insertion Map
Target$_A$-like
Target$_B$-like
Donor$_A$-like
Donor$_B$-like
Percent of reads
Genome coordinate (Mb)
Levenshtein distance

**D.**

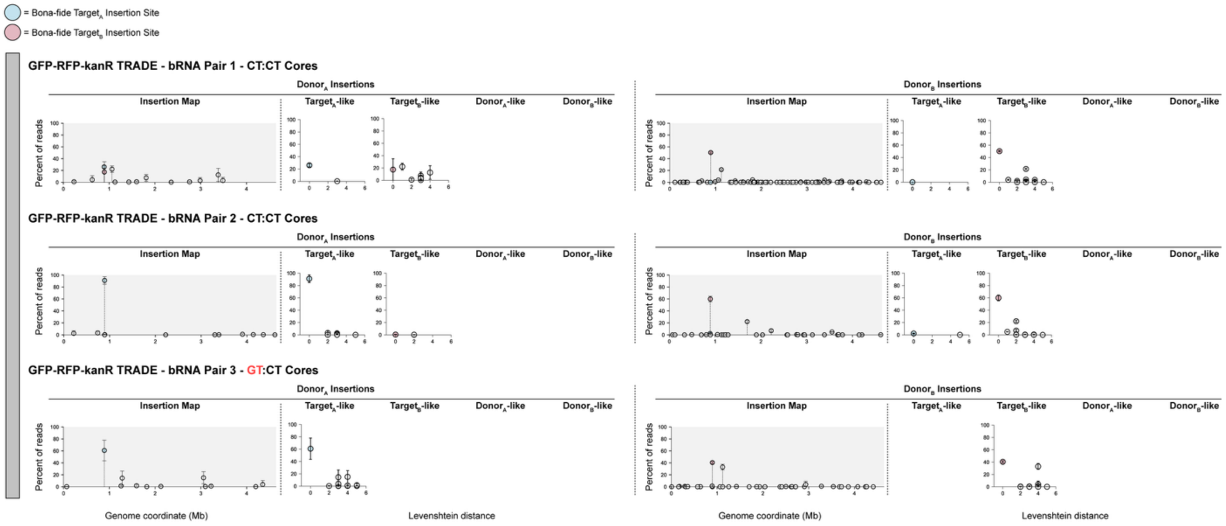
= Bona-fide Target$_A$ Insertion Site
= Bona-fide Target$_B$ Insertion Site
GFP-RFP-kanR TRADE - bRNA Pair 1 - CT:CT Cores
GFP-RFP-kanR TRADE - bRNA Pair 2 - CT:CT Cores
GFP-RFP-kanR TRADE - bRNA Pair 3 - GT:CT Cores
Donor$_A$ Insertions
Donor$_B$ Insertions
Insertion Map
Target$_A$-like
Target$_B$-like
Donor$_A$-like
Donor$_B$-like
Percent of reads
Genome coordinate (Mb)
Levenshtein distance

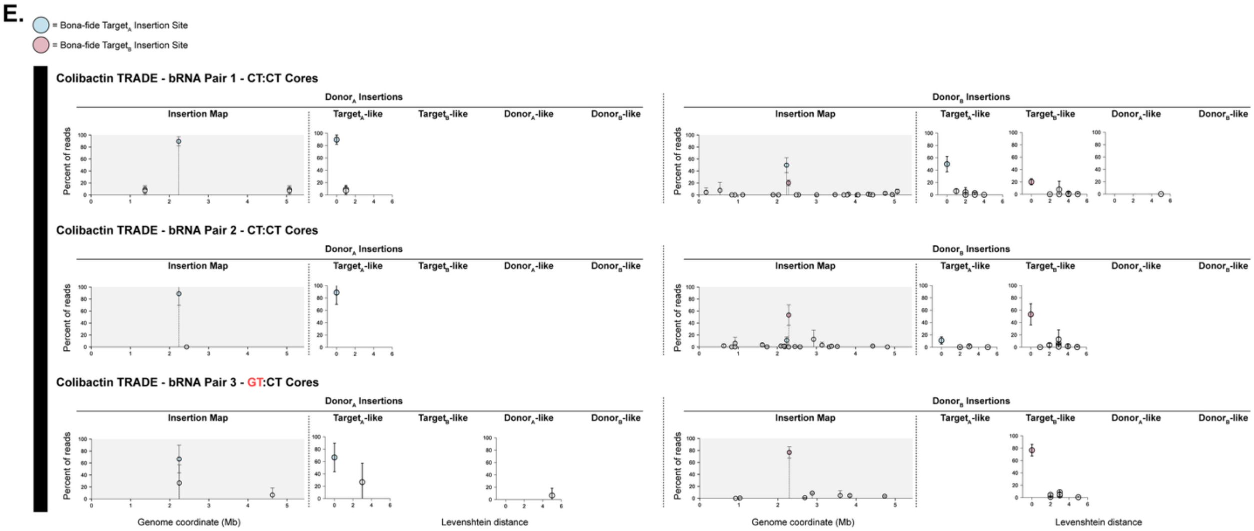


**Fig. S16. Dual bRNAs result in cross-reactivity that is mitigated by using orthogonal core sequences (Related to Fig. 4). A and B.** Following transformation and IS621 induction for TRADE editing of the *sfGFP-mRFP-kanR* and colibactin loci, respectively, AP-PCR and deep sequencing are used to map the insertion sites of both donor$_A$ and donor$_B$. Observed insertion sites are classified by sequence similarity as true programmed target$_A$, true programmed target$_B$, or off-target sites resembling target$_A$, target$_B$, donor$_A$, or donor$_B$. Cross-targets are defined as insertion of donor$_A$ into any *B-like* category and vice-versa. The fraction of reads appearing in each category are shown as a heat map, with cross-targets highlighted as a bar graph for each tested bRNA pair. **C, D, and E.** For each experiment utilizing dual bRNAs, genomewide integration sites of donor A and donor B are plotted as insertion maps. Additionally the abundance of each observed integration site is plotted as a function of Levenshtein distance to target$_A$, target$_B$, donor$_A$, or donor$_B$.

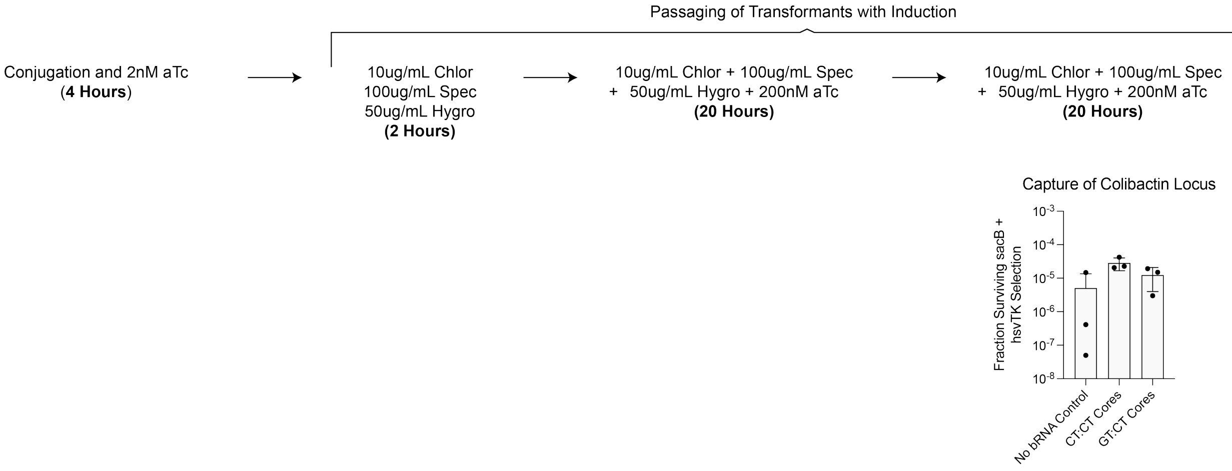


**Fig. S17. Selection methodology and raw selection efficiencies for capture of Colibactin locus with dual bRNAs (Related to Fig 5).** For the capture and translocation of the colibactin biosynthetic locus from the chromosome to a plasmid, transformation, induction, and selection protocols are outlined. For each bRNA pair, fractions surviving the respective selections are plotted.

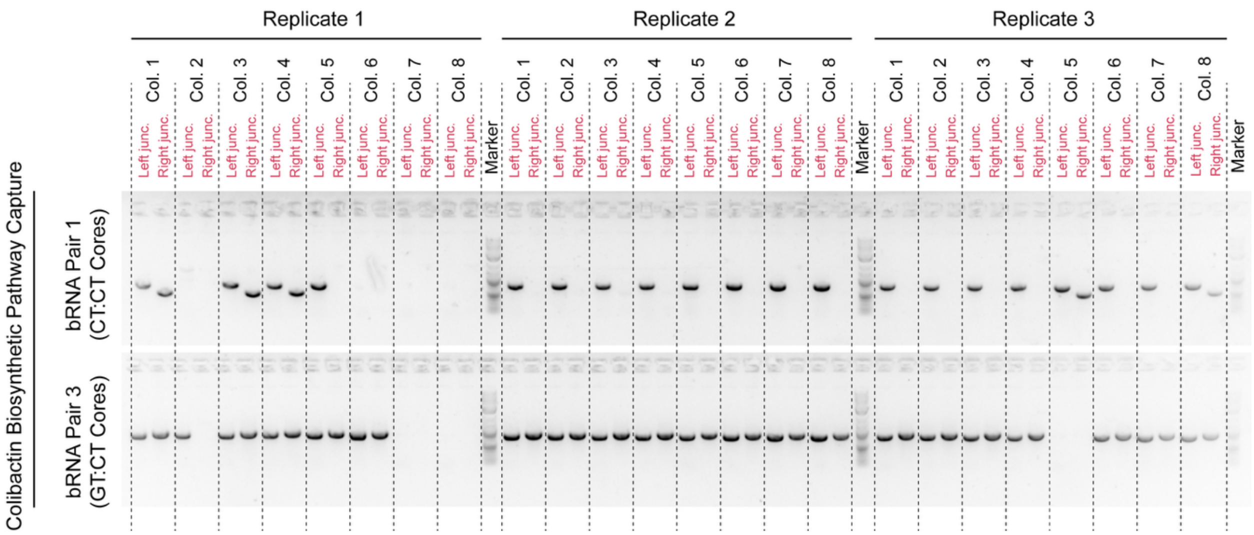


**Fig. S18. PCR-based quantification of the capture of the Colibactin locus with dual bRNAs (Related to Fig. 5).** Post pathway capture reaction and selection, individual colonies were screened via PCR to confirm both the left and right junctions between the captured colibactin biosynthetic pathway and the capturing vector.

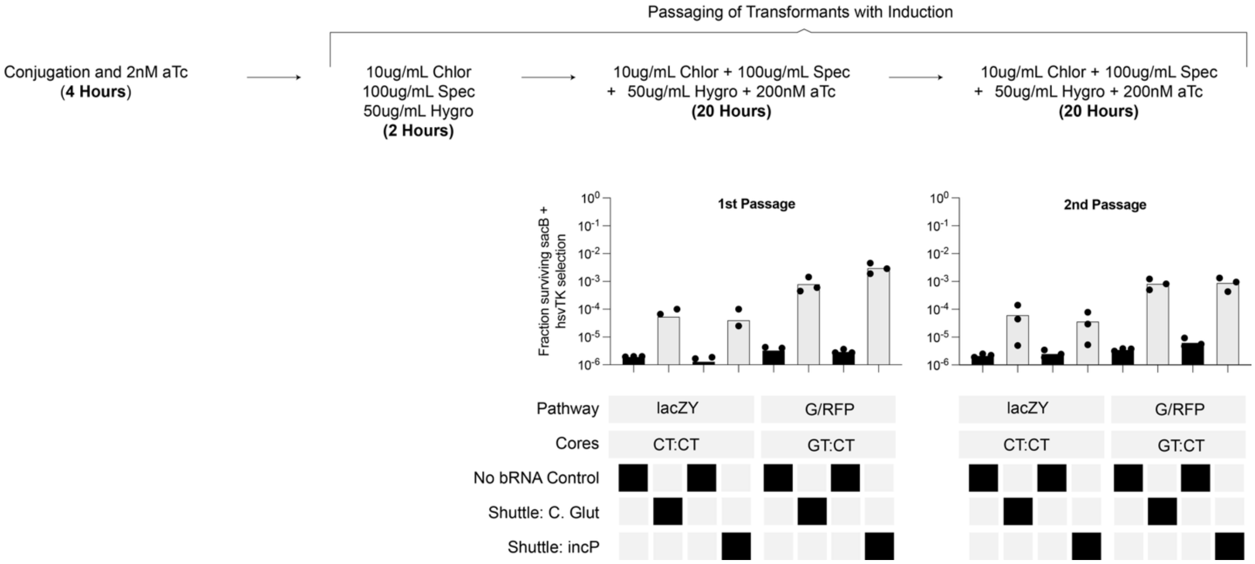


**Fig. S19. Selection methodology and raw selection efficiencies for capture of *sfGFP-mRFP-kanR* and *lacZY* loci with dual bRNAs (Related to Fig 5).** For the capture and translocation of the *lacZY* and *sfGFP-mRFP-kanR* biosynthetic loci from the chromosome to a shuttle plasmid, transformation, induction, and selection protocols are outlined. For each bRNA pair, fractions surviving the respective selections are plotted.

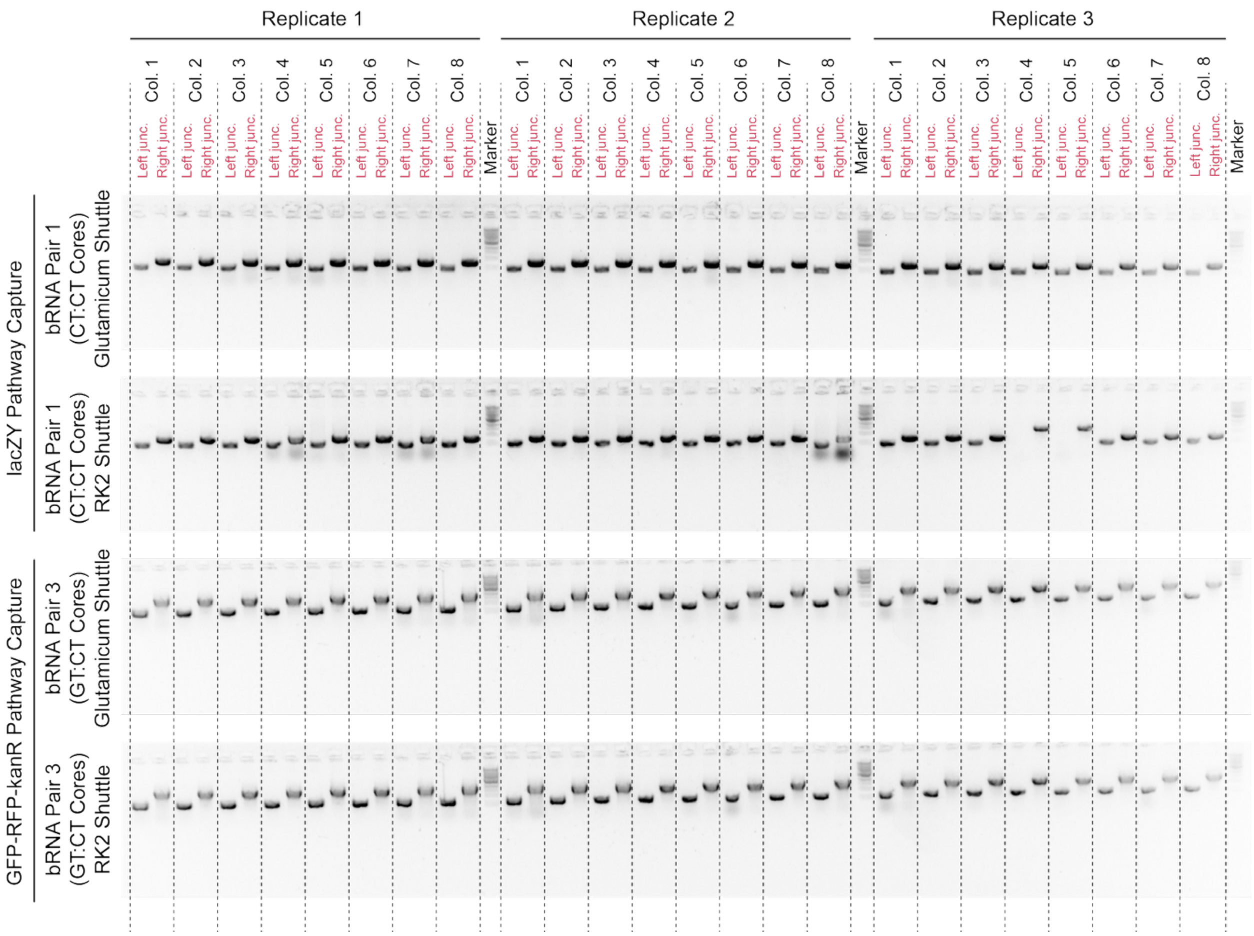


**Fig. S20. PCR-based quantification of the capture of the *sfGFP-mRFP-kanR* and *lacZY* loci with dual bRNAs (Related to Fig. 5).** Post pathway capture reaction and selection, individual colonies were screened via PCR to confirm both the left and right junctions between the captured pathway and the capturing shuttle vector.

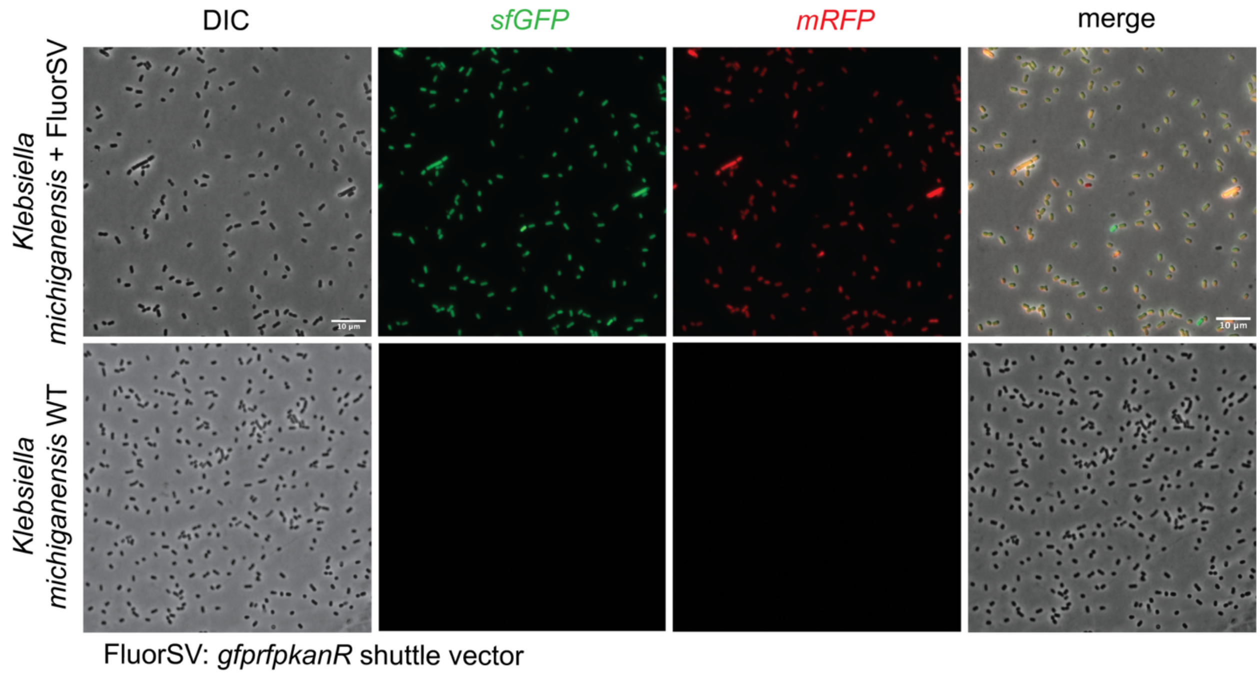


**Fig. S21. Fluorescence microscopy of *K. michiganensis* pCapture transformants (Related to Fig 5).** Following whole-plasmid sequencing verification of pCapture in *K. michiganensis*, low-melting-point agarose pad slides were made of the wild-type and the transconjugant strain. pCapture encoded the *sfGFP-mRFP-kanR* pathway captured from *Qi* coli and was stably maintained in overnight cultures of sequence-verified clones. Fluorescence microscopy validated the transfer of the GFP+ RFP+ phenotype in *K. michiganensis* transconjugants.

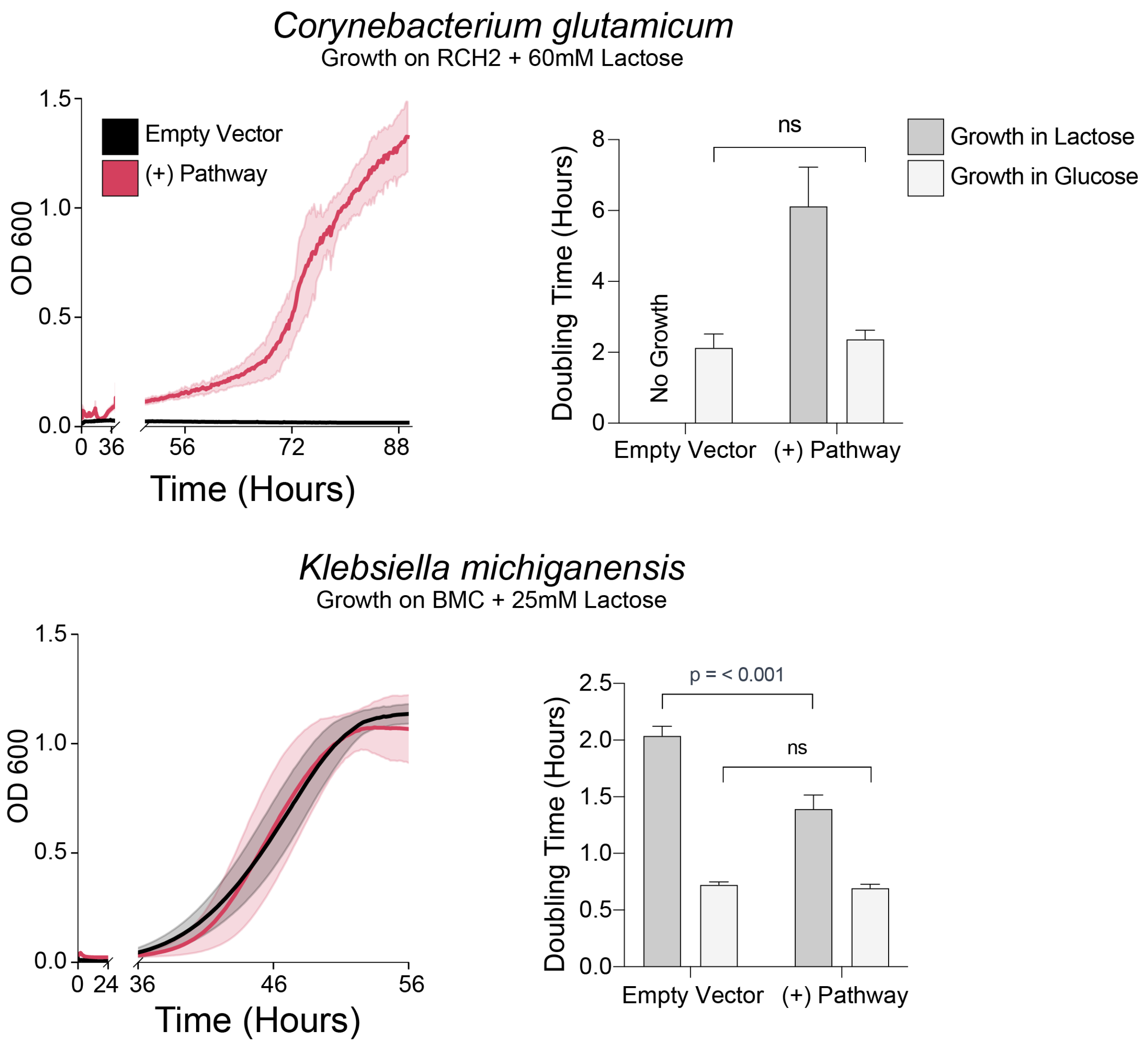


**Fig. S22. Transfer of *lacZY* pathway from *E. coli* to *C. glutamicum* and *K. michiganensis* enables and/or improves the ability to metabolize lactose as a sole carbon source (Related to Fig. 5).** Following conjugative transfer of the captured *lacZY* pathway from *E. coli* to *C. glutamicum* and *K. michiganensis*, kinetic growth in minimal media with lactose or glucose as a sole carbon source is monitored via OD600 absorbance in a plate reader. *C. glutamicum* gains the ability to grow on lactose as a sole carbon source, while *K. michiganensis* (already capable of lactose assimilation) achieves a growth rate enhancement. Growth curves are background-subtracted and fitted to a logistic curve to calculate the specific growth rates and doubling times in various conditions. A one-tailed t-test is performed to evaluate differences in doubling times.

**Table S1. (separate file)**

Strains, Plasmids, and Primers used in each experiment and for analysis are noted, with metadata where applicable.

**Table S2. (separate file)**

All bRNAs used in the study are listed, along with sequences and pertinent metadata

**Table S3. (separate file)**

Specific bRNAs that were used in each figure panel are noted, using consistent nomenclature

**Table S4. (separate file)**

For each of the 96 clones selected for analysis from the genomewide rearrangement assay, information enabling their mapping and classification into inversions and excisions is provided.

**Table S5. (separate file)**

For Broad Host Range Editing, pEdit vectors and culture conditions for each species are noted.

**Table S6. (separate file)**

For broad host range editing, AP-PCR derived pEdit insertion sites are tabulated along with absolute and relative sequencing read counts, classification into target-like and donor-like sequence, and Levenshtein distance to the respective classification.

**Table S7. (separate file)**

For broad host range editing, observed insertion frequency at 16S *rRNA* is tabulated using AP-PCR or colony PCR (of 11 CFUs per replicate) as the quantification method. Number of "target-like" sequences in the respective genomes are notes, categorized by Levenstein distance from the programmed target sequence.

**Table S8. (separate file)**

AP-PCR derived insertion sites of each donor in the dual bRNA assays is tabulated, along with pertinent metadata and abundance quantification

**Table S9. (separate file)**

For all deep sequencing analysis, the associated sequence fastQ file, its location, the corresponding reference genome, and pEdit plasmid are tabulated

**Data S1. (separate file)**

This file contains the tabulated data behind Figure 1.

**Data S2. (separate file)**

This file contains the tabulated data behind Figure 2.

**Data S3. (separate file)**
This file contains the tabulated data behind Figure 3.

**Data S4. (separate file)**
This file contains the tabulated data behind Figure 4.

**Data S5. (separate file)**
This file contains the tabulated data behind Figure 5.
